# Fundamental limits on dynamic inference from single cell snapshots

**DOI:** 10.1101/170118

**Authors:** Caleb Weinreb, Samuel Wolock, Betsabeh K. Tusi, Merav Socolovsky, Allon M. Klein

## Abstract

Single cell expression profiling reveals the molecular states of individual cells with unprecedented detail. However, because these methods destroy cells in the process of analysis, they cannot measure how gene expression changes over time. But some information on dynamics is present in the data: the continuum of molecular states in the population can reflect the trajectory of a typical cell. Many methods for extracting single cell dynamics from population data have been proposed. However, all such attempts face a common limitation: for any measured distribution of cell states, there are multiple dynamics that could give rise to it, and by extension, multiple possibilities for underlying mechanisms of gene regulation. Here, we describe the aspects of gene expression dynamics that cannot be inferred from a static snapshot alone and identify assumptions necessary to constrain a unique solution for cell dynamics from static snapshots. We translate these constraints into a practical algorithmic approach, Population Balance Analysis (PBA), which makes use of a method from spectral graph theory to solve a class of high dimensional differential equations. We use simulations to show the strengths and limitations of PBA, and then apply it to single-cell profiles of hematopoietic progenitor cells (HPCs). Cell state predictions from this analysis agree with HPC fate assays reported in several papers over the past two decades. By highlighting the fundamental limits on dynamic inference faced by any method, our framework provides a rigorous basis for dynamic interpretation of a gene expression continuum and clarifies best experimental designs for trajectory reconstruction from static snapshot measurements.

**Significance:** Seeing a snapshot of individuals at different stages of a process can reveal what the process would look like for a single individual over time. Biologists apply this principle to infer temporal sequences of gene expression states in cells from measurements made at a single moment in time. However, these inferences are fundamentally under-determined. Using a conservation law, we enumerate reasons that there is no unique dynamics associated with a single snapshot, limiting our ability to infer gene regulatory mechanisms. We then propose a method for dynamic inference that provides a unique dynamic solution under defined approximations and apply it to data from bone marrow stem cells. Overall, this study introduces formal biophysical approaches to single cell bioinformatics.

**Classification:** BIOLOGICAL SCIENCES / Systems Biology

## Introduction

Over the past few years, technologies for making genome-scale high-dimensional measurements on single cells have transformed our ability to discover the constituent cell states of tissues(1). These measurements enable a molecular dissection of biological tissues at the single cell level, across development, differentiation, disease onset, or in response to external stimuli. The most mature of these technologies, single cell RNA sequencing (scRNA-Seq), can be applied at relatively low cost to thousands and even tens of thousands of cells to generate an ‘atlas’ of cell states in tissues, while also revealing transcriptional gene sets that define these states(2, 3). Rapidly maturing technologies are also enabling single cell measurements of the epigenome(4), the proteome(5, 6), and the spatial organization of chromatin(7).

A more ambitious goal of single cell analysis is to describe dynamic cell behaviors, and by extension, to reveal dynamic gene regulation. Since high-dimensional single cell measurements are destructive to cells, they reveal only static snapshots of cell state. However, it has been appreciated that dynamic progressions of cell state can be indirectly inferred from population snapshots by methods that fit a curve or a tree to the continuous distribution of cells in high dimensional state space. To date, a number of methods have been published to address the problem of ‘trajectory reconstruction’ from single cell data. These methods have ordered events in cell differentiation(8-12), cell cycle(13), regeneration, and perturbation response(14). The most advanced algorithms have addressed increasingly complex cell-state topologies including branching trajectories(15).

Unfortunately, all attempts to infer dynamics from static snapshots of cell state face a common limitation: for any measured distribution of cells in high dimensional state space, there are multiple dynamics that could give rise to it, and by extension, multiple possibilities for underlying mechanisms of gene regulation. These limitations can apply even when sampling from multiple time points. Put differently, *any* computational method that reports a definite prediction for cell-state dynamics has made one choice among many about how to order observed cell states, whether or not the choice is made explicitly. To our knowledge, existing approaches rely on heuristic algorithms that do not explicitly state how bioinformatic decisions impact descriptions of biological dynamics. As such, the best methods for dynamic inference might be more accurately described as methods for non-linear dimensionality reduction, or ‘manifold discovery’: they robustly solve the problem of how to describe a static continuum of cell states using a small number of coordinates (often described as ‘pseudo-time’ coordinates), but they provide minimal guidance on how the observed static continuum (or ‘pseudo-time’) should be interpreted with respect to the many redundant dynamic processes that could give rise to it. Therefore, what assumptions must be made in dynamic inference – once the important task of manifold discovery is completed – remains an unsolved problem.

The difference between describing a manifold and describing its underlying biological dynamics becomes clear when considering the types of predictions one might make from data. Heuristic algorithms may be sufficient to provide an intuition for the biology and to identify dynamic genes and gene sets. However, quantitative predictions about cell behavior may require stronger forms of dynamic inference. We would be curious to know, for example, the relationship of real time to progression in ‘pseudo-time’ – the discrepancy between which has recently been explored(16) – and how to think about cell dynamics in the absence of a clear linear or branching structure. In systems where a cell can differentiate into multiple lineages, single cell data might encode how transcriptional programs influence fate bias; but it is not yet clear how to extract such information in a principled manner. The ambiguity of the biological dynamics associated with manifold descriptions also impacts the ability to infer regulatory mechanisms, since statements about mechanism necessarily entail specific hypotheses about cell trajectories. It follows that the limits on dynamic inference from single cell snapshots also affect attempts to reverse engineer gene regulatory networks(11) or to define “landscapes”(10) that confine cell dynamics in gene expression space.

Here we explore whether one can derive a framework for inferring cell state dynamics from static snapshots that overcomes the above ambiguities by identifying the critical assumptions implicit in inference, and identifying key fitting parameters that cannot be inferred from single cell data alone. With many algorithms now available for trajectory reconstruction from single cell data, our first focus is to define the limits of identifiability faced by *any* algorithm.

The second focus of this paper is to develop a practical algorithm for dynamic inference, which we call Population Balance Analysis (PBA). At one level, PBA provides a continuum description of cell states, just as existing methods do. However, PBA differs from existing algorithms in that it formally solves a problem of dynamic inference from biophysical principles, and can thus be considered predictive of cell dynamics under clearly stated assumptions. For example, it assigns to each transcriptional state a set of testable fate probabilities. We apply PBA to scRNA-seq data of hematopoietic progenitor cells (HPCs), reconciling these data with fate assays made over the past few decades in this system. Validation of novel PBA predictions in HPCs forms the subject of a second paper (Tusi, Wolock, Weinreb et al., *in submission*).

The biophysical foundation of PBA is embodied by a diffusion-drift equation over high-dimensional space, which, though simple to define, cannot be practically solved using established computational tools. We therefore invoke a novel, asymptotically exact and highly efficient solution to diffusion-drift equations using recent innovations in spectral graph theory. The ubiquity of diffusion-drift equations in fields of quantitative biology, physics and chemistry suggest that applications of these methods may exist in other fields. Overcoming this computational challenge represents the major technical contribution of this work.

## Results

### A first-principles relationship of cell dynamics to static observations

When reconstructing a dynamic process from single cell snapshot data, cells are typically observed in a continuous spectrum of states owing to asynchrony in their dynamics. The goal is to reconstruct a set of rules governing possible dynamic trajectories in high-dimensional space that are compatible with the observed distribution of cell states. The inferred rules could represent a single curve or branching process in gene expression space, or they could reflect a more probabilistic view of gene expression dynamics. In some cases, multiple time points can be collected to add clarity to the temporal ordering of events. In other cases, a single time point could capture all stages of a dynamic process, such as in steady-state adult tissue turnover.

To develop a framework for dynamic reconstruction from first principles, we wish to identify a general, model-independent, mathematical formulation linking cell dynamics to static observations. One possible starting point is the *population balance equation* (also known as the *flux balance law(17)*), which has the form:

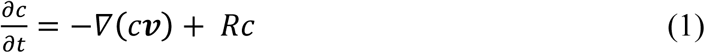

This partial differential equation provides a useful starting point for analysis, because it fully describes how the density of cells at a point in gene expression space depends on the average speed and direction of travel of cells. Formally, Eq. (1) states that in each small region of gene expression space, the rate of change in the number of cells (left-hand side of the equation) equals the net cell flux into and out of the region (right-hand side) (Fig. 1A). The equation introduces the cell density, *c*(*x,t*), which is the distribution of cell states from which we sample a static snapshot of cells in an experiment. This density depends on the net average velocity, ***v***(*x*), of the cells at point *x*, a feature of the dynamics that we wish to infer. Notably, being an average quantity, ***v*** is not necessarily a description of the dynamics of any individual cell, but it alone governs the form of the sampled cell density *c*. Eq. (1) also introduces a third variable: *R*(*x*) is a rate of cell accumulation and loss at point *x* caused by the discrete phenomena of cell proliferation and cell death, and by entrance and exit from the tissue being isolated for analysis. Though Eq. (1) is likely a good starting point for analyzing many biological systems, it nonetheless introduces some specific assumptions about the nature of cell state space. First, it approximates cell state attributes as continuous variables, though they may in fact represent discrete counts of molecules such as mRNAs or proteins. Second, it assumes that changes in cell state attributes are continuous in time,. This means, for example, that the sudden appearance or disappearance of many biomolecules at once cannot be described in this framework.

**Figure 1:**
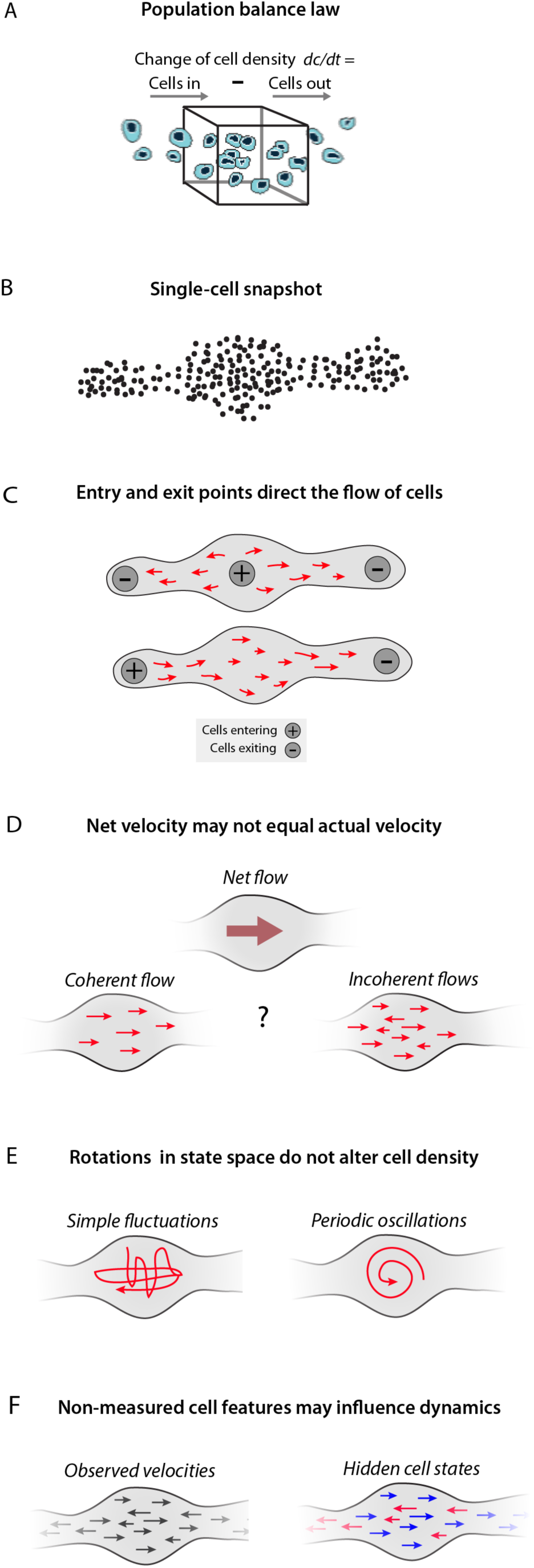
Symmetries and inhomogeneities of the Population Balance law set fundamental limits on dynamic inference. (A) Schematic of the population balance law [Eq. (1)], which serves as a starting point for inferring cell dynamics from high-dimensional snapshots. In each small region of gene expression space, the rate of change in cell density equals the net cell flux into and out of the region. Symmetries and unknown variables of the population balance law mean that there is no unique solution for dynamics from a static snapshot (B), shown schematically in (C-E). (C) Alternative assumptions on cell entry and exit rates across gene expression space lead to different dynamic solutions. (D) Snapshot data constrain only net cell flows through the population balance law, and not the noise in dynamic trajectories of individual cells. (E) A gauge symmetry of the population balance law means that static snapshots arising from periodic oscillations of cell state can also be explained by simple fluctuations that do not have a consistent direction and periodicity. (F) Hidden but stable properties of a cell – such as epigenetic state – allow for a superposition of cell populations following different dynamic laws. These unknowns are constrained by assumptions in any algorithm inferring dynamics from static snapshot data.

### Multiple dynamic trajectories can generate the same high dimensional population snapshots

Given knowledge of the cell population density, *c*(*x,t*), we hope to infer the underlying dynamics of cells by solving for the average velocity field ***v*** in Eq. (1). This approach falls short, however, because ***v*** is not fully determined by Eq. (1), and even if it were, knowing the average velocity of cells still leaves some ambiguity in the specific trajectories of individual cells. This raises the question: does there exist a set of reasonable assumptions that constrain the dynamics to a unique solution? To explore this question, we enumerate the causes of non-uniqueness in cell state dynamics, using a cartoon to introduce each cause (detailed in Figure 1), as well as referring to their mathematical foundation in Eq. (1).

1. *Assumed cell entry and exit points strongly influence inferred dynamics:* For the same data, making different assumptions about the rates and location of cell entry and exit lead to fundamentally different inferences of the direction of cell progression in gene expression space, as illustrated in Figure 1C. Cells can enter a system by proliferation, by physically migrating into the tissue that is being analyzed, or more mundanely by up-regulating selection markers used for sample purification (e.g. cell surface marker expression). Similarly, cells exit observation by cell death, physical migration out of the tissue being studied, or by down-regulation of cell selection markers. These events could be associated with particular gene expression states, or could occur broadly. Referring to Eq. (1), this discussion is formally reflected in the need to assume a particular form for the rate field *R*(*x*) when inferring dynamics ***v*** from the observed cell density *c.*
2. *Net velocity does not equal actual velocity:* A second unknown is the stochasticity in cell state dynamics, reflected in the degree to which cells in the same molecular state will follow different paths going forward. A net flow in gene expression space could result from imbalanced flows in many directions or from a single coherent flow in one direction (see Figure 1D). If the goal of trajectory analysis is to go beyond a description of what states exist and make predictions about the future behavior of cells (e.g. fate biases) given their current state, then it is necessary to account for the degree of such incoherence of dynamics. Incoherent dynamics also change inferences that might be made about underlying gene regulatory networks. Referring to Eq. (1), the net velocity field ***v*** reflects only the mean cell behavior, with individual cells deviating from the mean owing to stochastic gene expression or fluctuations in unmeasured quantities such as environmental cues.
3. *Rotations and oscillations in state space do not alter cell density:* Static snapshot data cannot distinguish periodic oscillations of cell state from simple fluctuations that do not have a consistent direction and periodicity (Figure 1E). As with incoherent motion above, predictive models may need to explicitly consider oscillatory behaviors. The inability to detect oscillations from snapshot data is formally reflected in Eq. (1) by invariance of the concentration *c* to the addition to ***v*** of arbitrary rotational velocity fields ***u*** satisfying ▽(c**u**) = 0.
4. *Hidden features of cell state can lead to a superposition of different dynamic processes:* Stable properties of cell state that are invisible to single cell expression measurements, such as chromatin state or tissue location, could nonetheless impact cell fate over multiple cell state transitions (Figure 1F). The existence of such long-term “hidden variables” would clearly compromise attempts to predict the future fate of a cell from its current gene expression state. Previously published algorithms for trajectory inference do not consider long-term hidden variables. This choice is inescapable for any modeling approach based on single cell RNA-seq or mass-cytometry data, since these measurement modalities simply do not capture every feature of a cell’s molecular state or its environment.

Because of these issues, no unique solution exists for dynamic inference. However, sensible predictions about dynamics can still be made by making certain assumptions. Our framework for cell trajectory analysis (see below) is based on explicit, reasonable assumptions that together are necessary and sufficient to constrain a unique solution (see Figure 2). These assumptions may nevertheless be inaccurate in certain situations.

### Construction of the Population Balance Analysis framework

To infer cell dynamics from an observed cell density *c*, we make the following assumptions.

#### The Fokker-Plank equation models memory-less cell state dynamics

The first assumption is that there are no hidden variables, meaning the properties of the cell available for measurement (such as its mRNA content) fully encode a probability distribution over its possible future states. This assumption is made implicitly by all current approaches to trajectory analysis and cell fate prediction, and we reflect on its plausibility in the discussion.

An equivalent statement of this first assumption is that cell trajectories are memory-less with respect to their measured properties, i.e. past states of the cell do not affect its future states other than through having led to its present state. If so, Eq. (1) can be treated as modeling a single dynamic process, described by a single local velocity field ***v***(*x*) subject to random, memoryless, noise. Eq. (1) in this case can be equivalently considered a continuum approximation of the chemical master equation (CME) that specifies the discrete stochastic molecular interactions underlying gene regulatory networks in the cell (18). Cell trajectories are modeled as biased random walks, with a deterministic component that reflects the reproducible aspects of cell state changes such as their differentiation through stereotypical sequences of states, and a stochastic component that reflects random fluctuations in cell state, partly driven by bursty gene-expression, fluctuations in cellular environment, and intrinsic noise from low molecular number processes. In turn, we can approximate Eq. (1) by a Fokker-Plank formalism, in which noise is treated as Gaussian in nature.

Fokker-Plank equations, which represent special cases of the Population Balance equation [Eq. (1)], have been applied previously to low dimensional biological processes, such as differentiation with a handful of genes(19) or a one-parameter model of cell cycle progression(13). Here, we apply them to high-dimensional data. Although Fokker-Plank descriptions are necessarily approximations, their emergence from first-principles descriptions of transcriptional dynamics(18), and their ubiquity in describing chemical reaction systems(20), justify their use instead of the more general form of Eq. (1). Specifically, the Generalized Fokker-Plank approximation takes the form of Eq. (1) with velocity field, 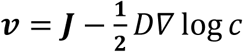, where the first term is a deterministic average velocity field, and the second term is a stochastic component of the velocity that follows Fickian diffusion with a diffusion matrix *D* (Figure 2). We assume here that *D* is isotropic and invariant across gene expression space. Though more complex forms of diffusion could better reflect reality, we propose that this simplification for *D* is sufficient to gain predictive power from single cell data in the absence of specific data to constrain it otherwise. The resulting Population Balance equation is thus,

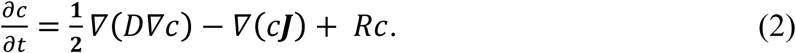

Eq. (2) explains the rate of change of cell density (∂*c*/∂*t*) as a sum of three processes: (1) stochastic gene expression, 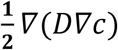, which causes cells to diffuse out of high-density regions in gene expression space; (2) convergences (and divergences) of the mean velocity field, ∇(*c****J***), which cause cells to accumulate (or escape) from certain gene expression states over time; and (3) as before, cell entry and exit rates, *R c*, will cause certain cell states to gain or lose cells over time.

#### Potential landscapes define a minimal model for dynamic inference

Our second assumption is that there are no oscillatory gene expression dynamics, which would appear as rotations in gene expression space. Though oscillations certainly do exist in reality – for example, the cell cycle – it is impossible to establish their existence from static snapshots alone. One is therefore forced to make an a priori assumptions about their existence, for example by specifically searching for signatures of a known oscillatory process in the data. For processes not known to be oscillatory, one can begin by making predictions of fate bias and temporal ordering while ignoring oscillatory phenomena. The utility of such predictions is supported by our analysis of single cell RNA-seq data in a later section.

In the Fokker-Plank formalism, the presumed absence of oscillations implies that the velocity field ***J*** is the gradient of a potential function *F* (i.e. ***J*** = - *∇F*). The potential would define a landscape in gene expression space, with cells flowing towards minima in the landscape, akin to energy landscapes in descriptions of physical systems. Applying the potential landscape assumptions to Eq. (2) gives rise to the simplified diffusion-drift equation below, where the potential is represented by a function *F*(*x*).

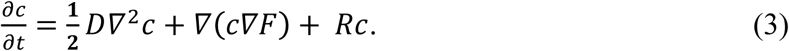

#### A recipe for dynamic inference from first principles

Equation (3) represents our best attempt to relate an observed density of cell states (*c*) to an underlying set of dynamical rules, now represented by a potential landscape (*F*) rather than the exact velocity field ***v***. Crucially, we have in these first few results sections: explained why the net cell velocity ***v*** is inherently unknowable; clarified why the description provided by a potential field *F* is the best that *any* method could propose without further knowledge about the system; and identified critical fitting parameters (*D*, *R* and ∂*c*/ ∂*t*), that are not revealed by single cell snapshot measurements, but are required for determining aspects of the dynamics such as temporal ordering of states and fate probabilities during differentiation. By starting from first principles, it becomes clear that these requirements are not limited to any particular algorithm; they affect any method one might develop for trajectory inference.

The challenge is now to develop a practical approach that relates the fitting parameters *D*, *R* and ∂*c*/ ∂*t* to dynamic predictions through Eq. (3). In the following, we focus on steady-state systems where ∂*c*/ ∂*t* = 0, and use prior literature to estimate *R*. We report results for a range of values of *D*. Building on the work here, more elaborate approaches could be taken, for example determining *R* from direct measurements of cell division and cell loss rates or integrating data from multiple time points to estimate ∂*c*/ ∂*t*, thus generalizing to non-steady-state systems.

### Reducing to practice: solving the Population Balance equation with spectral graph theory

Equipped with single cell measurements and estimates for each fitting parameter, we now face two practical problems in using of Eq. (3) to infer cell dynamics: the first is that Eq. (3) is generally high-dimensional (reflecting the number of independent gene programs acting in a cell), but numerical solvers cannot solve diffusion equations on more than perhaps ten dimensions. Indeed, until now, studies that used diffusion-drift equations such as Eq. (3) to model trajectories(10, 13, 19) were limited to one or two dimensions, far below the intrinsic dimensionality of typical single cell RNA-seq data(21). The second practical problem is that we do not in fact measure the cell density *c*: we only sample a finite number of cells from this density in an experiment.

Overcoming these problems represents the main technical contribution of this paper. We drew on a recent theorem by Ting, Huang and Jordan in spectral graph theory(22) to extend diffusion-drift modeling to arbitrarily high dimension. The core technical insight is that an asymptotically-exact solution to Eq. (3) can be calculated on a nearest-neighbor graph constructed with sampled cells as nodes, rather than on a low-dimensional sub-space of gene expression as performed previously (e.g.(13)). Our approach, which we call Population Balance Analysis (PBA) actually *improves* in accuracy as dimensionality increases, rewarding high-dimensional measurements. We thus avoid conclusions based on low-dimensional simplifications of data, which may introduce distortions into the analysis. In practice, some intermediate degree of dimensionality reduction could still be useful (say, to tens or hundreds of dimensions), a point elaborated in the Discussion. The supplement of this paper provides technical proofs and an efficient framework for PBA in any high-dimensional system.

**Figure 2:**
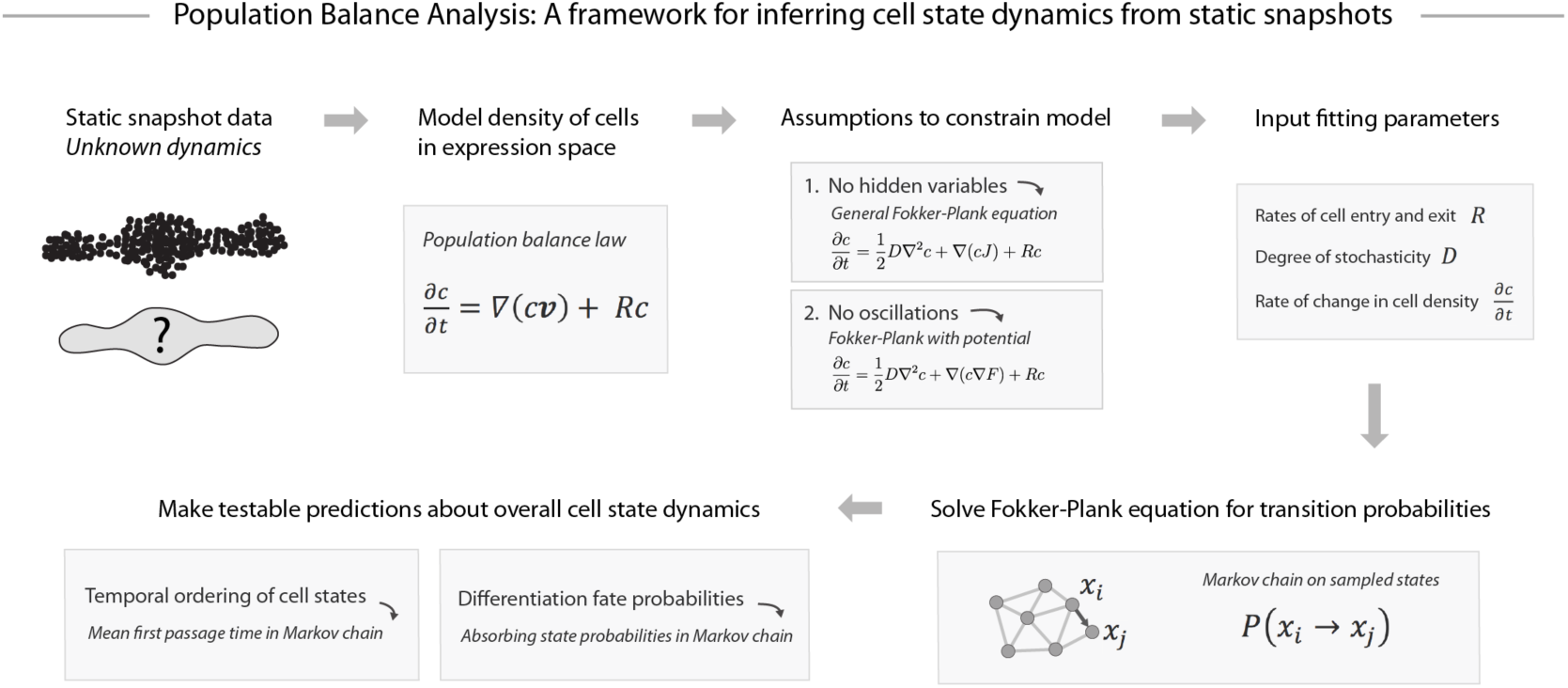
Population Balance Analysis. Though many dynamics are consistent with a given static snapshot of cell states, testable assumptions can constrain a unique solution. Shown schematically here is Population Balance Analysis (PBA), one such approach to dynamic inference under explicit assumptions. PBA constrains the population balance law by assuming: no hidden variables; a dynamics described by a potential landscape (see text for details); and fitting parameters that incorporate prior knowledge or can be directly measured. The resulting diffusion-drift equation is solved asymptotically exactly in high dimensions on single cell data through a graph theoretic result (supplement and ref(2 1)). The PBA algorithm outputs transition probabilities for each pair of observed states, which can then be used to compute dynamic properties such as temporal ordering and fate potential.

The inputs to PBA are a list of sampled cell states ***x*** = (*x*_1_, … *x*_*N*_), an estimate *R* = (*R*_1_, … *R*_*N*_) for the net rate of cell accumulation/loss at each state *x*_*i*_, and an estimate for the diffusion parameter *D*. We are assuming steady-state, so ∂*c*/ ∂*t* = 0. The output of PBA is a discrete probabilistic process, i.e. a Markov chain, that generates the Fokker-Planck equation (Eq. (3)), and thus describes the transition probabilities between the states *x*_*i*_. The analysis is asymptotically exact in the sense that – if a potential exists and the estimates for *R* and *D* are correct – then the inferred Markov chain will converge to the underlying continuous dynamical process in the limit of sampling many cells (*N* → ∞) (Theory supplement, Theorem 4).

PBA computes the transition probabilities of the Markov chain using a simple algorithm, which at its core involves a single matrix inversion. Briefly:

1. Construct a *k*-nearest-neighbor (knn) graph *G*, with one node at each position ***x***_*i*_ extending edges to the *k* nearest nodes in its local neighborhood. Calculate the graph Laplacian of *G*, denoted *L*.
2. Compute a potential *V* =L^+^ *R*, where L^+^ is the pseudo-inverse of *L*
3. To each edge (*x*_*i*_ → *x* _*j*_), assign the transition probability

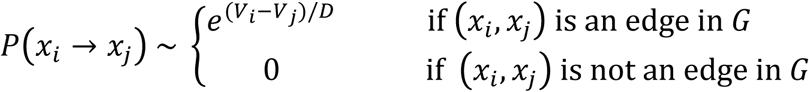

With the Markov chain generating Eq. (3) available, it is possible to calculate the temporal ordering of states (via mean first passage time), and the fate biases of progenitor cells in a differentiation process (via absorbing state probabilities), by integrating across many trajectories (see Figure 2). These calculations are simple, generally requiring a single matrix inversion. Specific formulas are provided in the Theory Supplement Section 3. Code for implementing these and other aspects of PBA is available online at https://github.com/AllonKleinLab/PBA.

**Figure 3:**
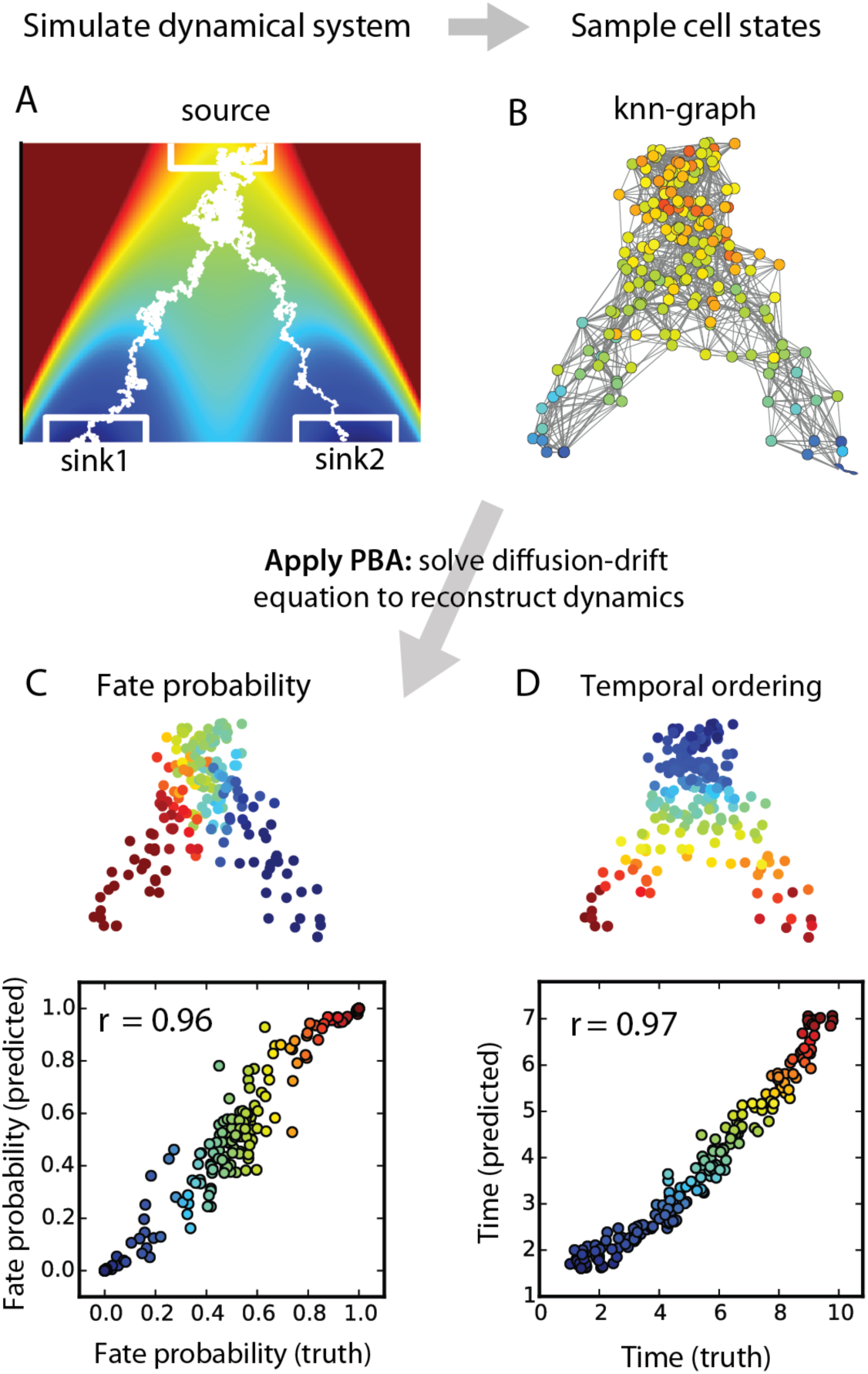
Demonstration of PBA on a simulated high-dimensional differentiation process. (A) Cells emerge from a proliferating bi-potent state (source) and differentiate into one of two fates (sinks 1 and 2) in a high-dimensional gene expression space, with two dimensions shown. Heat map colors show a potential field containing the cell trajectories. Example trajectories shown in white. (B) Static expression profiles sampled asynchronously through differentiation serve as the input to PBA, which reconstructs trajectories and accurately predicts future fate probabilities (C) and timing (D) of each cell.

### PBA accurately reconstructs dynamics of simulated differentiation processes

We tested PBA on a sequence of simulations, first using an explicit model of diffusion-drift process, and then moving on to direct simulations of gene regulatory networks. In the first simulation (Figure 3; Supplementary Figure 1), cells drift down a bifurcating potential landscape into two output lineages. Cell trajectories span a 50-dimensional gene expression space (two of which are shown in Fig. 3A). With 200 cells sampled from this simulated system (Figure 3B), PBA predicted cell fate probabilities and temporal ordering of the measured cells. PBA made very accurate predictions (Pearson correlation, *ρ > 0.96*, Fig. 1C-D) if provided with correct estimates of proliferation, loss and stochastisticity (parameters *R* and *D*). Estimates of temporal ordering remained accurate with even 5-fold error in these parameters (*ρ* > 0.93), but predictions of fate bias degraded (*ρ* > 0.77; Supplementary Figure 2A-D). Thus even very rough knowledge of the entry/exit points in gene expression space is sufficient to generate a reasonable and quantitative description of the dynamics. Interestingly, PBA also remained predictive in the presence of implanted oscillations (Supplementary Figure 3, fate probability *ρ* > 0.9; temporal ordering *ρ* > 0.8). In addition, the simulations confirmed the theoretical prediction that inference quality improves as the number of noisy genes (dimensions) increases, and as more cells are sampled: maximum accuracy in this simple case was reached after ∼100 cells and 20 dimensions (Supplementary Figure 2e-g). These simulations showcase the ability of PBA to not just *describe* continuum trajectories, but to additionally *predict* cell dynamics and by extension cell fate. At the same time, they show the fragility of dynamic inference to information not available from static snapshots alone.

**Figure 4:**
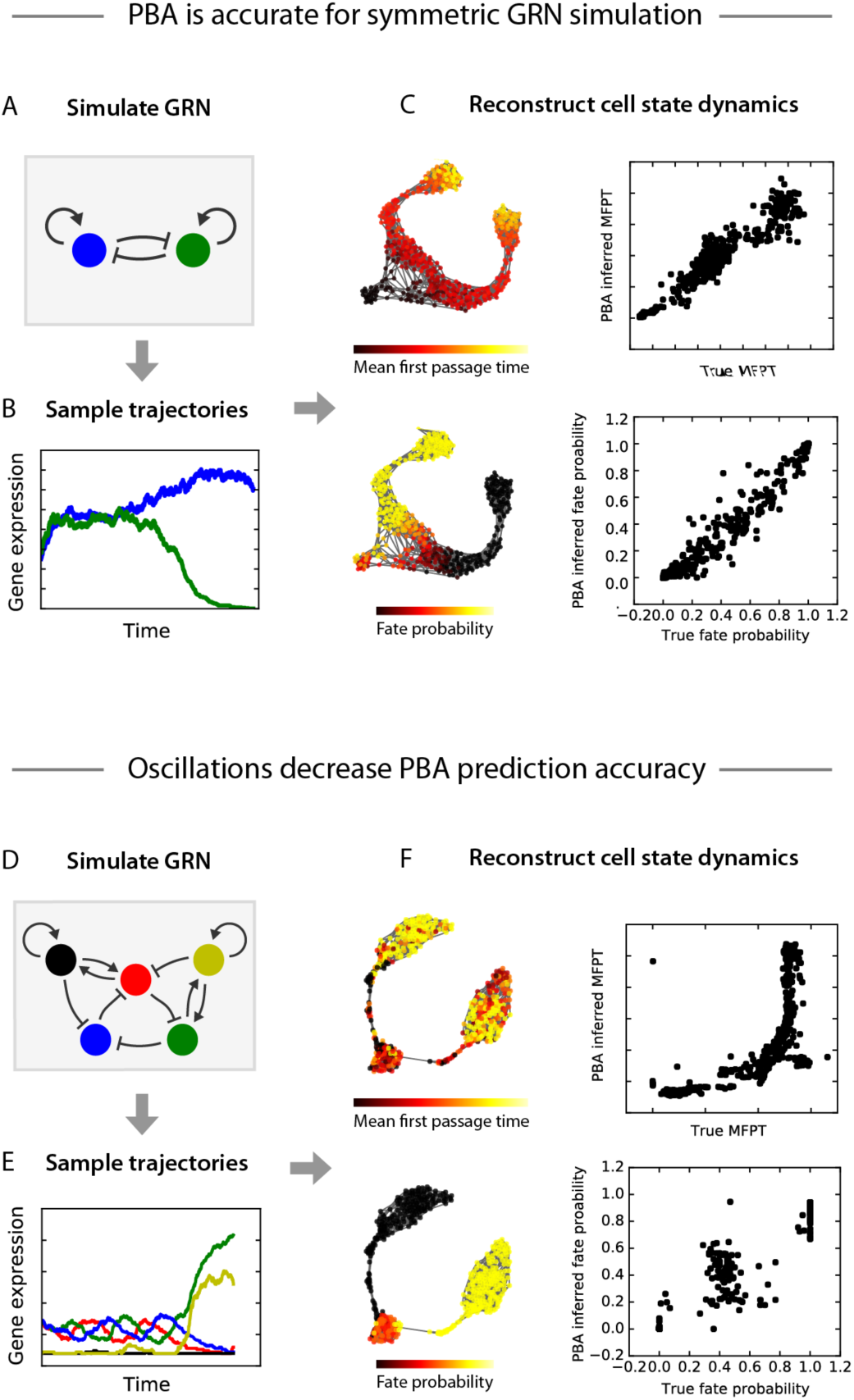
Test of PBA on cell states from a simulated gene regulatory network. (A) We tested PBA on cell states sampled from a gene regula tory network (GRN) composed of two genes that repress each other and activate themselves. (B) Trajectories from this GRN begin in an unstable state with both genes at an intermediate level and progress to a stable state with one gene dominating. (C) PBA was applied to a steady-state snapshot of cells from this process (shown on the left using a force-directed layout generated by SPRING). The resulting predictions for temporal ordering (top) and fate probability (bottom) are compared to ground-truth. (D) To challenge PBA, we defined a GRN with two stable states that compete with semi-stable limit cycle. (E) Trajectories from the GRN begin with all genes oscillating and then progress to stable state where one pair of genes dominates. (F) PBA was applied to a steady-state snapshot of cells from this process, with predictions for ordering (top) and fate proba bility (bottom) compared to ground-truth.

Having demonstrated the accuracy of PBA on an explicit model of a diffusion-drift process, we next tested its performance on gene expression dynamics arising from gene regulatory networks (GRNs) (Figure 4). As before, we simulated cell trajectories, obtained a static snapshot of cell states, and supplied PBA with this static snapshot as well as the parameter *R* encoding the location of entry and exit points. We began with a simple GRN representing a bi-stable switch, in which two genes repress each other and activate themselves (Figure 4A). Simulated trajectories from this GRN begin with both genes at an intermediate expression level, but quickly progress to a state where one gene dominates the other (Figure 4B). In addition to the two genes of the GRN, we included 48 uncorrelated noisy dimensions. With 500 cells sampled from this process, PBA predicted cell fate bias and temporal ordering very well (r>0.98 for fate bias and r > 0.89 for ordering; Figure 4C), though the precise accuracy depended on the assumed level of diffusion *D* (Supplementary Figure 4).

PBA assumes the absence of oscillations in gene expression space. Therefore, it is unclear how well PBA can infer cell trajectories that result from GRNs with oscillatory dynamics. We simulated an oscillatory GRN in the form of a “repressilator” circuit(23) with the addition of positive feedback loops that create two “escape routes” leading to alternative stable fixed points of the dynamics (Figure 4D). Simulated trajectories from this GRN begin with all genes oscillating, followed by a stochastic exit from the oscillation when one of the genes surpasses a threshold level (Figure 4E). With 500 cells sampled from this process, PBA was significantly less accurate than for the previous simulations (Figure 4F). Though PBA correctly identified which cells were fully committed to the two ‘escape routes’, it was entirely unable to resolve the fate biases of cells in the uncommitted oscillatory state. PBA also made poor predictions of mean first passage time, underestimating the amount of time that cells spent in the oscillatory state. Unsurprisingly, when the assumptions of PBA are strongly violated its prediction accuracy suffers.

### PBA predictions of fate bias in hematopoiesis reconcile past experiments

To test PBA on experimental data from real biological systems, we made use of single cell gene expression measurements of 3,803 adult mouse hematopoietic progenitor cells (HPCs) from another study by our groups (Tusi, Wolock, Weinreb et al., *in submission;* data at https://www.ncbi.nlm.nih.gov/geo/query/acc.cgi?token=mrqdcsemnxatnev&acc=GSE89754).

HPCs reside in the bone marrow and participate in the steady-state production of blood and immune cells through a balance of self-renewal and multi-lineage differentiation. Fate commitment of HPCs is thought to occur through a series of hierarchical fate choices, investigated over the past four decades through live cell tracking, in vitro colony-forming assays and transplantation of defined sub-populations of HPCs(24). Depictions of the HPC hierarchy invoke a tree structure, with gradual lineage-restriction at branch points. However the precise tree remains controversial(25, 26), since existing measurements of fate potential reflect a patchwork of defined HPC subsets that may have internal heterogeneity(27) and provide only incomplete coverage of the full HPC pool. We asked whether PBA applied to single cell RNA profiling of HPCs could generate predictions consistent with experimental data, and possibly help resolve these controversies by providing a global map of approximate cell-fate biases of HPCs.

The single cell expression measurements – derived from mouse bone marrow cells expressing the progenitor marker Kit – represent a mixture of multipotent progenitors as well as cells expressing lineage commitment markers at various stages of maturity. Since PBA prescribes analysis of a k-nearest-neighbor (knn) graph of the cells, we developed an interactive knn visualization tool for single cell data exploration, called SPRING, (kleintools.hms.harvard.edu/tools/spring.html; (28)). The SPRING plot (Figure 5A) revealed a continuum of gene expression states that pinches off at different points to form several downstream lineages. Known marker genes (Supplementary Table 1) identified the graph endpoints as monocytic (Mo), granulocytic (G), dendritic (D), lymphoid (Ly), megakaryocytic (Mk), erythroid (Er) and basophil (Ba) progenitors (Supplementary Figure 5); we also identified cells in the graph expressing HSC markers. The lengths of the branches reflect the timing of Kit down-regulation and the abundance of each lineage.

**Table 1:**
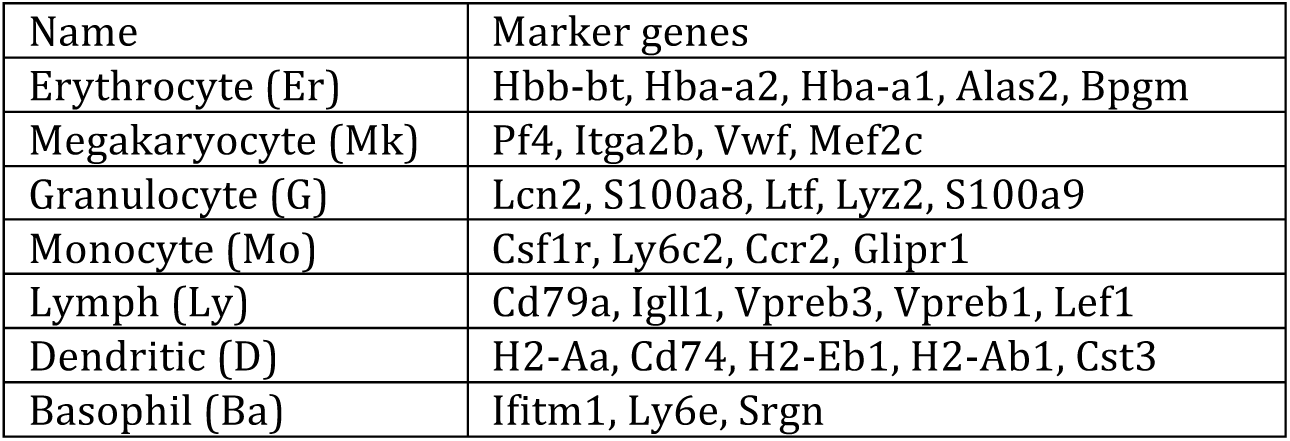
Marker genes used to identify the most mature cells in each lineage.

For steady-state systems, PBA requires as fitting parameters an estimate of the diffusion strength *D*, and the net rates of cell entry and exit at each gene expression state (*R*). We estimated *R* using prior literature (see methods), and tested a range of values of *D*. All results that follow hold over the physiological range of PBA parameter values (Supplementary Figure 6).

We compared PBA results to previously reported fate probabilities by localizing reported cell types on the graph using published microarray profiles. Remarkably, for a panel of twelve progenitor cell populations from six previous papers(29-34) (Supplementary Table 2) the PBA predicted fate outcomes (Figure 5B) closely matched fate probabilities measured in functional assays (defined as the proportion of clonogenic colonies containing a given terminal cell-type; see Supplementary Figure 5). The main qualitative disagreement between PBA predictions and experiment was in the behavior of Lin^-^Sca1^-^ Kit^+^IL7R^-^FcgR^low^CD34^-^ HPCs, previously defined as megakaryocyte-erythroid precursors (MEP)(29). Our prediction was that these cells should lack megakaryocyte potential, which is indeed consistent with recent studies(25, 27, 35). Excluding these cells, our predicted fate probabilities matched experimental data with correlation ρ=0.91 (Fig. 5C). In a second paper (Tusi, Wolock, Weinreb et al., *in submission*), we test several novel predictions in hematopoiesis emerging from PBA.

**Table 2.** Summary of 12 HPC subpopulations with microarray profiles and fate assays from previous papers, used to validate PBA

\*NM = not measured
| Name | Lineage Potential<sup>1</sup> (%)<br>Er-Mk-G-Mo-Ly-D | Microarray reference | Microarray accession codes (replicates) | Differentiation assay | Differentiation reference | Figure in original reference | Num. cells<sup>2</sup> |
| --- | --- | --- | --- | --- | --- | --- | --- |
| PreMegE | 50-50-0-0-NM-NM | Pronk, 2007 | GSM2076(82-86) | Agar | Pronk, 2007 | Figure 3 | 12<sup>3</sup> |
| PreCFU-E | 95-5-0-0-NM-NM | Pronk, 2007 | GSM2076(90-91) | Agar | Pronk, 2007 | Figure 3 | 12<sup>3</sup> |
| MkP | 0-100-0-0-NM-NM | Pronk, 2007 | GSM2076(87-89) | Agar | Pronk, 2007 | Figure 3 | 12<sup>3</sup> |
| Pre GM | 12-8-45-35-NM-NM | Pronk, 2007 | GSM2076(79-81) | Agar | Pronk, 2007 | Figure 3 | 12<sup>3</sup> |
| CMP | 14-16-37-32-NM-NM | Teng, 2008 | GSM7911(17-18) | Methylcellulose | Akashi, 2000 | Figure 1 | 12<sup>4</sup> |
| MEP | 45-55-0-0-NM-NM | Teng, 2008 | GSM7911(08-09) | Methylcellulose | Akashi, 2000 | Figure 1 | 30<sup>4</sup> |
| GMP | 0-0-60-40-NM-NM | Teng, 2008 | GSM7911(19-21) | Methylcellulose | Akashi, 2000 | Figure 1 | 18<sup>4</sup> |
| CDP | NM-NM-0-0-0-100 | Teng, 2008 | GSM7911(14-16) | In vivo | Naik, 2007 | Supp Fig 1 | 3<sup>3</sup> |
| CLP | 0-0-0-0-100-0 | Teng, 2008 | GSM5383(48-50) | In vivo | Rumfelt, 2006 | Figure 2 | 3<sup>4</sup> |
| MDP | 0-0-0-75-0-25 | Teng, 2008 | GSM7911(05-07) | S17 stroma | Fogg, 2006 | Figure 2B | 12<sup>4</sup> |
| MPP2 | 25-25-25-25-NM-NM | Pietras, 2015 | GSM16746(29-30) | Methylcellulose | Pietras, 2015 | Figure 2 | 3<sup>5</sup> |
| MPP3 | 10-10-40-40-NM-NM | Pietras, 2015 | GSM16746(31-33) | Methylcellulose | Pietras, 2015 | Figure 2 | 3<sup>5</sup> |

<sup>1</sup> Lineage potential refers to the proportion of colonies/mice that produce a terminal cell-type when inoculated with the given progenitor population. Potentials are normalized to add up to one. When measurements were made for HSCs, the potentials were renormalized so that HSC would have uniform potential across cell types.
<sup>2</sup> All cell numbers represent a ratio with respect to short-term stem cells (ST-HSC). When data was not available for a specific progenitor population, we used data from a population with the same functional potential, or otherwise made a conservative guess.
<sup>3</sup> Conservative guess
<sup>4</sup> (Busch, 2015)
<sup>5</sup> (Pietras, 2015)

**Figure 5:**
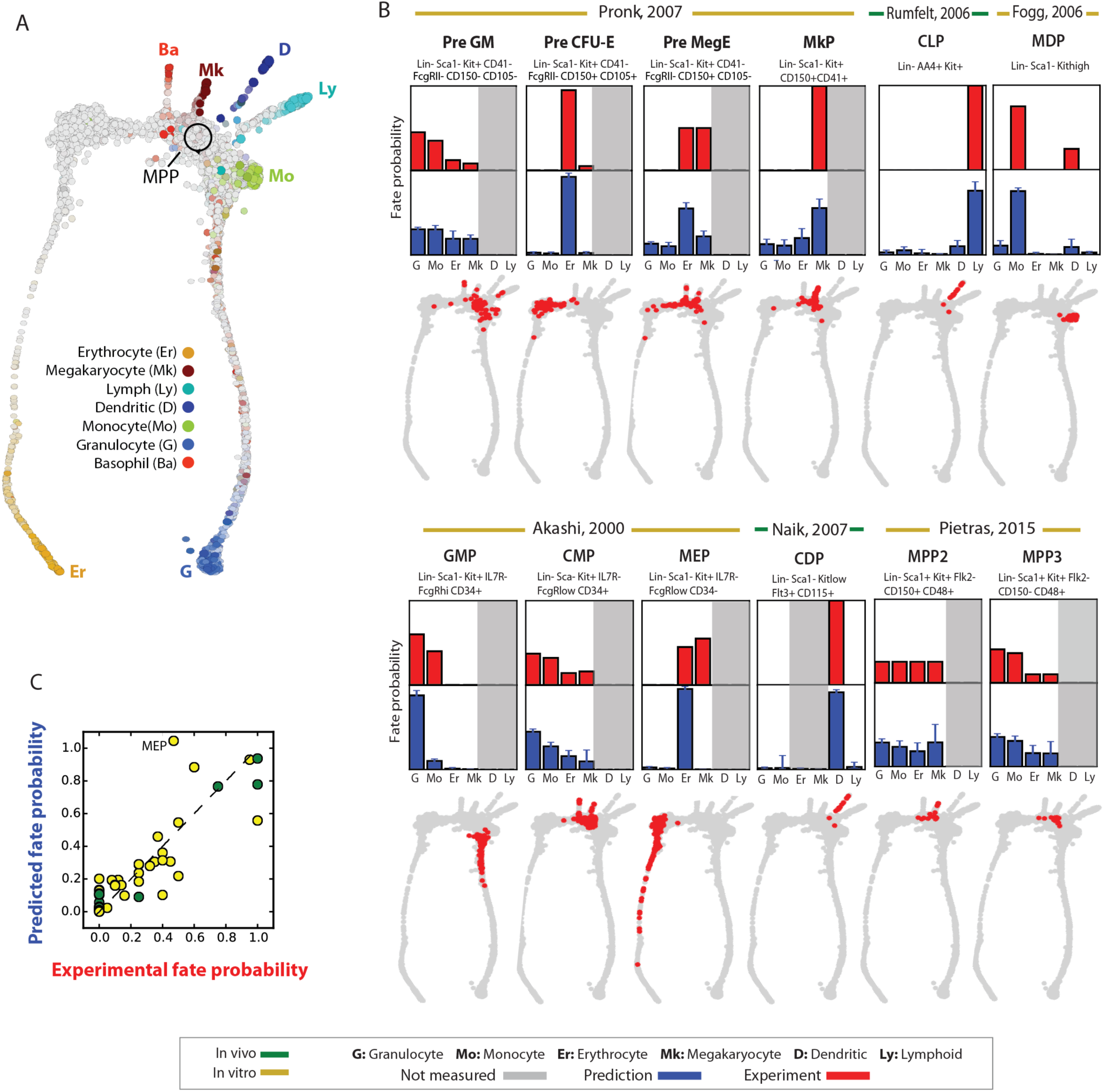
Population BaIance reproduces known fate probabilities of hematopoietic progenitor cell (HPC) subpopulations from single cell data. (A) Single-cell profiles of 3,803 Kit+ HPCs reveal a continuum of gene expression states that pinches off at different points to form seven downstream lineages. Cells in this map are colored by marker gene expression and are laid out as a k-nearest-neighbor (knn) graph using SPRING, an interactive force-directed layout software. (B) PBA is applied to HPCs, and the predicted fate probabilities (blue bars) are compared to those observed experimentally (red bars) for reported HPC subpopulations (red dots on gray HPC map; identified using transcriptional similarity to existing microarray profiles for each reported subpopulation). Cell fates predicted by PBA but not measured experimentally are shaded gray. Error bars = 90% confidence intervals across 120 parameter combinations for the PBA pipeline. (C) Summary of comparisons made in (B); green points = in vivo measurements; yellow points in vitro measurements.

## Discussion

In developing PBA, we hoped that an algorithm with clear assumptions would help to clarify the ways in which data analysis might mislead us about the underlying biology. More practically, we hoped that the algorithm would suggest how to best design experiments to extract dynamic information from static measurements, and how to visualize single cell data to preserve aspects of the true dynamics. We discuss a number of points that follow from our analysis, along with a note about the technical underpinnings of PBA.

### Experimental design for trajectory reconstruction from static snapshot measurements

We have shown that accurate dynamic inference requires knowledge of the density of cells in high dimensional state space, as well as the rates of cell entry and exit across the density. These requirements immediately suggest a set of principles for experimental design to optimize dynamic inference. First, to minimize distortions in the cell density in gene expression space it is useful to profile a single, broad population, and to avoid merging data from multiple subpopulations fractionated in advance. Second, if cells of interest are sorted prior to analysis, it is best to minimize the number of sorting gates and enrichment steps, since each introduces an additional term to the entry/exit rates and subsequently a risk of distortion to the inferred dynamics. The HPC dataset analyzed in this paper was well suited for trajectory reconstruction because it consisted of a single population, enriched using a single marker (Kit). This contrasts with previous single-cell RNA seq datasets of hematopoietic progenitors that included a composite of many sub-populations(36) or used complex FACS gates to exclude early progenitors(27).

### Experimental methods for beating the limits of trajectory reconstruction

Given the inherent limits of trajectory reconstruction from single cell snapshots alone, orthogonal experimental data is required to disambiguate the true dynamics. In differentiation systems, pulse-chase experiments – where cells labeled in a given state are followed over time – could be used to infer rates of cell entry and exit by reporting on the flux of cells into different lineages. A previous study(37) quantified the transition rates between different FACS-defined hematopoietic compartments using a Tie2 – driven reporter to pulse label HSCs; the same assay could be coupled to single-cell sequencing to enable direct fitting of the entry/exit parameter *R*. Clonal barcoding is another approach that would powerfully complement dynamical inference algorithms such as PBA: the dispersion of clones in high dimensional state space should constrain the stochasticity in the dynamics, allowing estimates of the diffusion constant *D* or even allowing consideration of non-uniform diffusion across gene expression space. Finally, live imaging of one or a few reporters could provide information on the significance of oscillatory behaviors that are not detectable in snapshot data.

**Figure 6:**
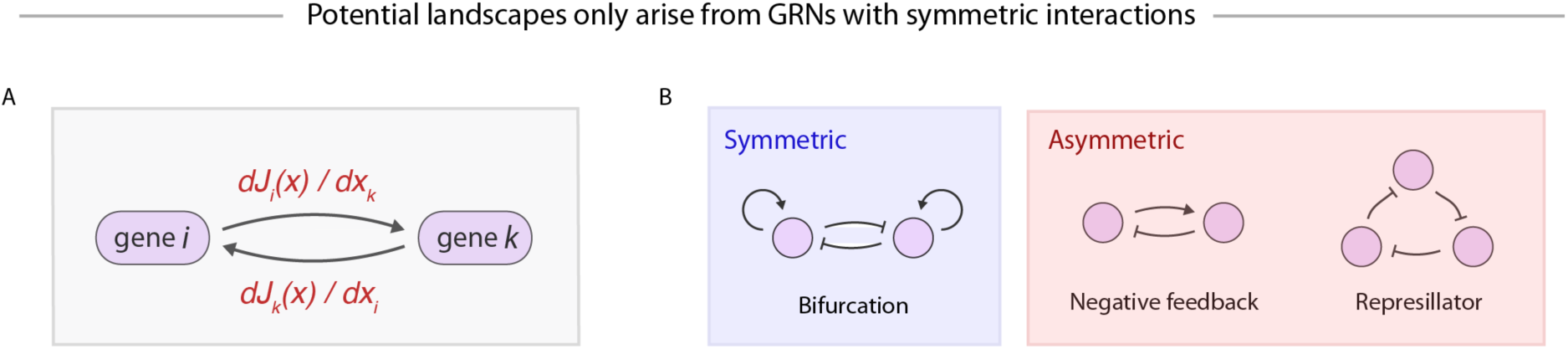
Potential landscapes arise from symmetric GRNs. (A) Inferences about the deterministic component of average cell velocities, J(x), can be interpreted as statements about an underlying gene regulatory network (GRN), with dJi/dxj giving the sensitivity of the dynamics of gene ito the expression level of gene j. (B) The existence of a potential landscape-driven dynamics implies that the underlying GRN has strictly symmetric interactions, which allows for some common gene regulatory motifs but rules out many others.

### How Population Balance Analysis could go wrong

To constrain a unique solution for trajectory reconstruction, PBA makes several strong assumptions, such as the absence of hidden variables and the absence of oscillations in gene expression space. Our simulations show that PBA is highly accurate for systems that meet these assumptions, but incorrectly infers dynamics for systems that break them. The agreement of prediction to fate commitment assays when we applied PBA to single cell profiles of hematopoietic progenitor cells suggests that, despite some sensitivity to assumptions, accurate inference is possible for complex differentiation systems. However, in cases where oscillatory dynamics strongly influence cell fate, or where hidden variables play a large role, single cell snapshot data could be misleading, and methods that infer dynamics from continua of cell states, such as PBA, may be ill suited.

In general, the impact of hidden variables on cell state dynamics remains unclear. Though there are many stable and possibly unobserved properties that impact a cell’s behavior – including chromatin state; post-translational modifications; cellular localization of proteins; metabolic state; and cellular micro-environment – it is possible that these properties percolate to some aspect of cell state that is observed, e.g. effecting a change in the expression of at least one gene measured by RNA-seq. By altering the observed state, such variables would thus not be hidden. For example, chromatin state exists in constant dialogue with transcriptional state, and is well reflected in mRNA content.

### Normalization, Principal Components Analysis, and other coordinate transformations

In this study, we described a framework for modeling the movement of cells in a space of gene expression, the units of which might be considered to be (dimensionless) counts of individual molecules. How then should one think about routine transformations of gene expression coordinates performed during practical low-level processing of single-cell expression data, such as transformation into logarithmic space, or dimensionality reduction by principal components analysis (PCA)? Here the asymptotic analysis of PBA makes clear that coordinate transformations may not be important when cells are densely sampled, as they should leave the empirical single cell graph topology unchanged. The equations of PBA are indeed invariant to coordinate transformations, with the exception of the diffusion operator, which is isotropic and spatially homogeneous but may not remain so upon coordinate system transformation. Since our assumption of isotropic and invariant diffusion is already an approximation, it does not support *a priori* one coordinate system over another. For small and noisy data sets, the choice of coordinate system could affect conclusions however, and it is probably best to use the coordinate system that provides the richest view of single-cell population structure, or that agrees most with known biology.

### Fundamental limits on the inference of gene regulatory networks

One promise of single cell expression measurements is their possible use for reconstructing gene regulatory networks (GRNs) (2, 11). However, since any GRN model entails specific hypotheses about the gene expression trajectories of cells, efforts to infer GRNs from single cell data must also confront the limits of knowledge identified in our framework. In particular, GRN inference may benefit from an explicit consideration of cell entry and exit rates (embodied by *R*) and the rate of change in the cell density (∂*c* / ∂*t*), as well as acknowledging the inability to distinguish oscillations from fluctuations.

Indeed, the inability to detect oscillations in single cell data, embodied in our framework by the use of a potential landscape, suggests severe limits on the types of underlying gene regulatory relationships that can be modeled. In fact, potential landscapes can only emerge from GRNs with strictly symmetric interactions, meaning every “arrow” between genes has an equal and opposite partner. This result follows from observing that the “arrows” in a GRN describe the influence of gene *i* on gene *j*, which is given by ∂***J***_i_ /∂*x*_*j*_ (Figure 6A), where ***J*** is the deterministic component of average cell velocities (see Equation 2). The assumption of a potential landscape (i.e. ***J*** = -∇*F*) then imposes symmetry on the GRN because ∂*J*_*i*_ ∂*x*_*j*_ = ∂*J*_*j*_ ∂*x*_*i*_=- ∂^2^ F/∂*x*_*i*_∂*x*_*j*_. Though a few well-known GRN motifs follow this symmetry rule – such as the “bistable switch” resulting from the mutual inhibition of two genes – many others do not, such as negative feedback loops and oscillators (Figure 6B). Potential landscapes are frequently invoked to explain gene expression dynamics(10, 38, 39), and we have shown them to be useful for predicting HPC fate outcomes in the context of PBA. It seems paradoxical that a tool that provides realistic phenomenological descriptions of gene expression dynamics reflects an entirely unrealistic picture for the underlying gene regulatory mechanisms. Resolving this paradox is an interesting direction for future work.

### How should we visualize single cell data?

At its core, the PBA algorithm performs dynamic inference by solving a diffusion-drift equation in high dimensions. This computation relies on a 2011 result in spectral graph theory by Ting Huang and Jordan(22) that describes the limiting behavior of *k*-nearest-neighbor graph Laplacians on sampled point clouds. Interestingly, several recent studies(8, 40, 41) have developed k-nearest neighbor graph-based representations of single cell data, and others have suggested embedding cells in diffusion maps(21, 42) on the basis of other similarity kernels. It has been unclear, until now, how to evaluate which of these different methods provides the most useful description of cell dynamics. Our technical results (Theorems 1-4 in the supplement) confirm that certain graph representations provide an asymptotically exact description of the cell state manifold on which dynamics unfold, suggesting them to be useful techniques for visualizing single cell data sets. Therefore PBA formally links dynamical modeling to choices of single cell data visualization.

## METHODS

### 1. Population balance analysis (PBA) source code and inputs

The core functions of PBA are implemented in python scripts on our github page: https://github.com/AllonKleinLab/PBA. The github page contains example files sufficient to reproduce the main calculations in this paper. In the theoretical supplement, we develop the rigorous foundations for PBA, provide detailed pseudo-code for the PBA algorithm and prove mathematically that it is asymptotically exact when sufficient cells are sampled and when PBA assumptions (see main text) are satisfied.

PBA was applied to simulated datasets and to experimental data from hematopoietic progenitor cells (HPCs) by calling PBA subroutines as follows:

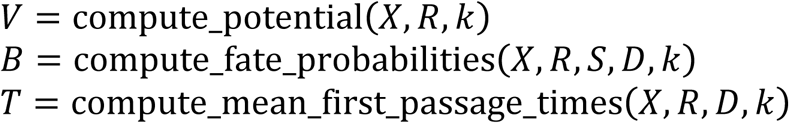

In each case, the inputs to PBA are: a collection of single-cell expression profiles *X*, where *X* _*i j*_ is an expression matrix of gene *j* in cell *i*; prior estimates of the relative rates of proliferation and loss provided at each sampled gene expression state as a vector *R* of length *n*; the exit rates of cells into *M* terminal fates specified as a matrix *S* of size (*n* x *M*); and a diffusion constant (*D*) that reflects stochasticity in the dynamics. The number of neighbors of the nearest neighbor graph, *k,* is a fitting parameter, but results are not sensitive to the choice of *k* (see Supp. Fig. 2b). We used a range of *k* values for all analyses. The first output of PBA is a vector giving the values of the potential *V* at the *n* sampled expression states. *V* is then used to calculate a set of transition probabilities between sampled cells, from which we further derive terminal fate probabilities of each sampled cell provided as a matrix *B* of size (*n* x *M*), as well as the conditional mean first passage time between every pair of sampled cells provided as a matrix T of size (*n* x *n*).

For the simulated data, parameters *R* and *D* are determined in Methods sections (2-5). For the experimental data, we fitted *R* and *D* in Methods sections (7-8). For the analysis of HPCs, we normalized and reduced dimensionality of the raw expression data to generate a reduced matrix *X* as input for PBA (see Methods section 6).

Note that in general, *R* and *D* are partially redundant, since multiplying both by a common factor does not change the fate probabilities output by PBA.

### 2 Simulating a diffusion-drift process (Figure 3)

Data used to test PBA were generated from a simulated differentiation process in which initially bipotent cells choose one of two fates. In each simulation a single cell is generated in a gene expression state chosen uniformly at random in an *m*-dimensional box (the entry point) corresponding to an initial gene expression profile, with boundaries −0.5<*x*_1_<0.5, 0.75<*x*_2_<1, and 0<*x*_k>2_<1 (Supp. Fig. 1). The cell gene expression profile is then updated over time using a Langevin simulation, meaning that a cell with initial position *x*(*t*) is given a new position as follows:

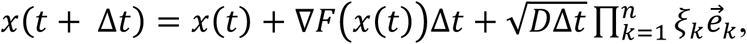

where ξ_1_, …, ξ _*m*_∼Gaussian(0,1) are independent random variables sampled from a normal distribution, 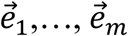 are unit vectors in each of the *m* dimensions, the simulation time step is Δ*t* = 0.001 and *D* = 0.05. The mean gene expression velocity for cells in state *x* is the gradient of the *m*-dimensional potential field *F*, which defines a bifurcation in two dimensions with a quadratic basin in the remaining *m* - *2* dimensions:

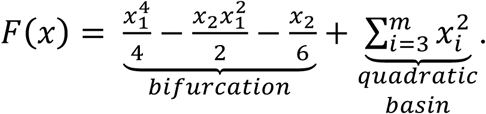

The simulation is terminated when a cell enters either of two box-shaped regions (the exit regions) where they were removed at rate *R* = 5 (boundaries shown in Supp. Fig. 1). This means that in a time step Δ*t*, cells in an exit box are removed with probability 1 – *e*^-RΔt^ ≈ 5Δ*t*. All simulations used *m*=50 dimensions, except for Supp. Fig. 2g, where *m* was varied from 2-50. The simulation was repeated to generate *N* cell trajectories, and each cell was then “sampled” at a time selected uniformly at random to generate mock single cell data set for PBA. All calculations were performed with *N*=200 sampled cells, except for Supp. Fig. 2e, where *N* was varied from 10-300.

For comparison of PBA predictions to the true dynamics (Figs. 1e-f and Supp. Figs. 2d-g), the “true fate probability” was defined for each sampled state *x*_*i*_ by carrying out a further 1000 Langevin simulations for each *x*_*i*_ as the initial condition, and recording the fraction of simulations terminating in exit box 2. The “true time since entry” was assigned to each *x*_*i*_ as the mean simulation time to reach *x*_*i*_ and its 5 nearest neighbors from a look-up table of 10,000 simulated trajectories.

### 3. Tests of PBA on simulated diffusion-drift process (Figure 3)

PBA was used to predict fate probability and time since entry for each sampled state *x*_*i*_. PBA takes as input the entire point cloud {*x*_*i*_}, as well as prior estimates of the entry/exit rates *R*_*i*_ at each point. For the main test of PBA (Figure 3) we used the true *R*_*i*_ values (*R*_*i*_ = 5 for cells in the entry-box; *R*_*i*_ = −5 for cells in the exit-boxes; *R*_*i*_ = 0 otherwise). For the robustness test in Supp. Fig. 2a-b we used false assumptions about entry/exit rates as indicated in the figure panels. For the robustness test in Supp. Fig. 2c-d we used false assumptions about the diffusion constant *D* as indicated in the figure panels. Changing *D*→*D’* is equivalent to scaling *R* uniformly, *R*→*R’=(D/D’)R*.

### 4. Testing the effect of gene oscillations on PBA predictions for diffusion-drift simulation (Supplementary Figure 3)

PBA assumes that the gene expression dynamics that give rise to a given set of sampled points {*x*_*i*_} is the gradient of a potential. However, this solution is not unique. The PBA solution implicitly assumes there are no rotations in gene expression space: a rotational field would not change the static density of cell states, and so it is invisible in a single cell sampling experiment. The effect of rotations on PBA predictions was tested by implanting rotational fields into the above simulations (Supp. Fig. 3). The rotational field used was

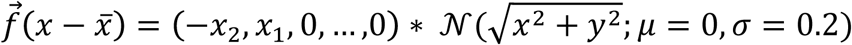

where 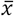 is the center-point of the rotational field and 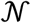 denotes a normal distribution. Langevin simulations were repeated as described above after adding this velocity field to the potential gradient velocity field. PBA predictions were repeated as described above to generate the results shown in Supp. Fig. 3.

### 5. Tests of PBA on simulated GRNs (Figure 4)

We used the Gillespie algorithm(1) to generate molecular counts for the simulations of gene regulatory networks (GRNs) in Figure 4. In every case, we supplemented the simulated counts with additional noisy dimensions (values drawn from a Gaussian), so that the total dimensionality of the data was always 50.

For the GRN in Figure 4A, we implemented the following stochastic chemical reactions (*x*_1_ represents the green node, *x*_2_ represents the blue node).

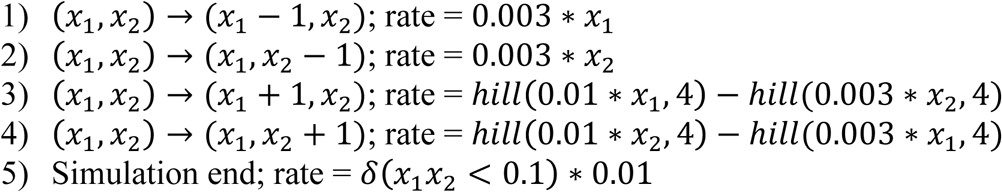

For the GRN in Figure 4D, we implemented the following stochastic chemical reactions, where the variables *x*_i_ correspond to the colors in the figure as follows (*x*_1_, red; *x*_2_, green; *x*_3_, blue; *x*_4_, black; *x*_5_, yellow)

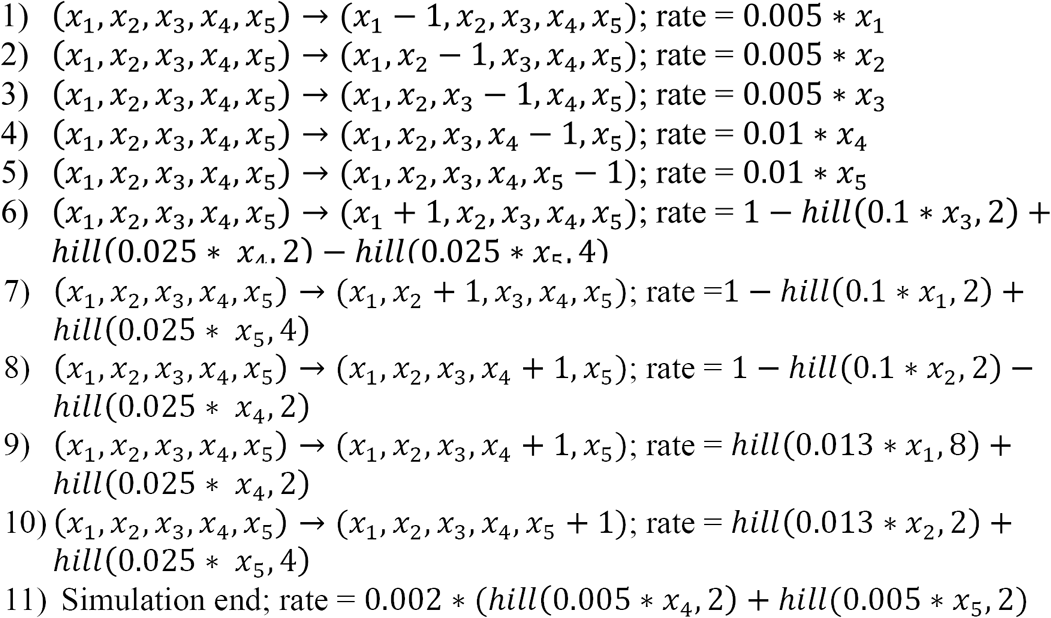

### 6. Data processing and normalization of single-cell RNA-seq data

Single-cell gene expression data from adult mouse bone marrow cells expressing Kit are reported and processed in another paper from our groups (Tusi, Wolock, Weinreb et al., *in submission*) Recapping in brief, reads were mapped as described in (2) to produce a (cell x gene) matrix of unique molecular identifier (UMI) counts that served as the starting point for the analysis in this paper.

Data was filtered to remove cells with < 1000 total UMIs. Visualization of the remaining cells in tSNE revealed three aberrant clusters of cells: one cluster strongly expressed mitochondrial genes and likely contained to stressed cells; the other two clusters co-expressed markers for distinct mature lineages (erythrocyte/macrophage and erythrocyte/granulocyte) and likely contained doublets. We removed all three aberrant clusters, resulting in 3803 cells.

Single cell data was then prepared for PBA by normalizing the total gene expression counts in each cell as described in (2). Genes with mean expression > 0.05 across the data set, and Fano Factor > 2, were then used to perform principal components analysis down to *p* dimensions, for *p* = 40, 50, 60, 70, 80, 90. When applying PBA, we also used a range of graph neighbor connectivities *k* (*k*=10 – 30). In Figure 5, we report medians and confidence intervals of fate probabilities for all 120 combinations of *p* and *k*.

### 7. Determining entry and exit parameters (R) for PBA analysis of HPCs

To apply PBA to hematopoietic differentiation, we estimated the entry/exit rates *R* from considerations of the proliferation rate and exit rates of Kit+ HPCs as follows. In adult hematopoiesis, all progenitors including HSCs express Kit, but eventually down-regulate it as they terminally differentiate. Thus, no cells enter the experimental system other than through proliferation of existing Kit+ HPCs, but there is a steady outflow (exit) owing to down-regulation of Kit as cells differentiate. We encoded this exit as negative *R* values for the top 10 cells with highest marker gene expression for each of the seven terminal lineages (Supplementary Figure 5 and Supplementary Table 1). We assigned different magnitudes of *R* for each of the seven lineages using a fitting procedure (see next paragraph). All remaining cells were assigned a uniform positive value of *R*, corresponding to a uniform proliferation rate, based on recent studies (3, 4) that found roughly similar growth rates across hematopoietic progenitor compartments. The magnitude of the growth rate was chosen so that ∑ *R*_*i*_ = 0, reflecting a steady state in the total number of cells.

The flux of cells down-regulating Kit for each lineage varies widely between different hematopoietic lineages. This impacts PBA because it directly sets the relative magnitude of *R* for each lineage, although the simulations indicate that predictions do not require very accurate flux estimates. Because the flux of Kit+ cells from each lineage is not generally known, we fitted the seven fluxes by requiring that PBA reproduce measured fate probabilities of hematopoietic stem cells (HSCs). We performed a separate fitting for each of the studies shown in Figure 5 (see Supplementary Figure 7). When a study did not report fate probabilities for HSCs, we assumed a uniform distribution. We identified HSCs in our data by comparison to a microarray profile of HSCs, as described in Methods section 9.

### 8. Determining the diffusion rate (D) for PBA analysis of HPCs

The diffusion rate (*D*) controls the level of stochasticity in the PBA model. The exact value of *D* cannot be directly measured, but it is possible to constrain *D* using known quantities. We defined a physiologically plausible range by scanning through different values of *D* and checking the number of PBA-predicted multipotent cells for each value (see Supp. Fig. 7). We used prior literature (see https://www.immgen.org/, (4) and Methods section 9) to estimate that 2-20% of Kit+ bone marrow cells are multipotent. We defined a cell in our dataset as multipotent (for a given value of *D*) if it satisfied *P*(fate) > 1/14 for all 7 fates.

### 9. Validation of PBA-predicted fate probabilities for HPC

To validate PBA predictions of HPC fate probability, we compared them to the fate probabilities of 12 HPC subsets measured in previous studies (Table 2; Supplementary Figure 6). For each of the 12 cell surface marker-defined hematopoietic compartments, we used a published microarray profile to search for similar cells in our own dataset using a naïve Bayesian classifier, implemented as follows.

The Bayesian classifier assigns cells to microarray profiles based on the Likelihood of each microarray profile for each cell, with the Likelihood calculated by assuming that individual mRNA molecules in each cell are multinomially sampled with the probability of each gene proportional to the microarray expression value for that gene. Consider a matrix *E* of mRNA counts (UMIs) with *n* rows (for cells) and *g* columns (for genes), and also a matrix *M* with *m* rows (for microarray profiles) and *g* columns for genes. *M* was quantile normalized and then each microarray profile was normalized to sum to one. *E* was previously normalized in Methods section 5. The (*n*×*m*) matrix *S*_*ij*_ giving the Likelihood of each microarray profile *j* for each cell *i* is,

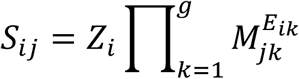

where *Z*_*i*_ is a normalization constant that ensures ∑_*i*_ *S*_*ij*_ = 1.

We assigned *N*_*j*_ cells with highest log-Likelihoods to each microarray profile *j*, with *N*_*j*_ determined from prior literature to reflect the abundance of each cell type among HPCs (see Supp. Table 2). Previous studies only provide abundance ratios between cell compartments, so we estimated *N*_*j*_ values by first estimating the number of ST-HSCs in our data, and then multiplying this value by the relative of abundance of each compartment compared to ST-HSCs. We estimated that the number of ST-HSCs in our data set was *N* = 5, reasoning that: (1) 1% of adult bone marrow is Kit+ (i.e. in our dataset); (2) the proportion of HSCs in adult bone marrow is 1-2 in 100,000 (5) and thus 1-2 in every 1,000 Kit+ cells is an ST-HSC; (3) our dataset contains approximately 5000 cells. Final assignments are indicated on the knn graphs in Figure 5.

---

**Supplementary Figure 1:**
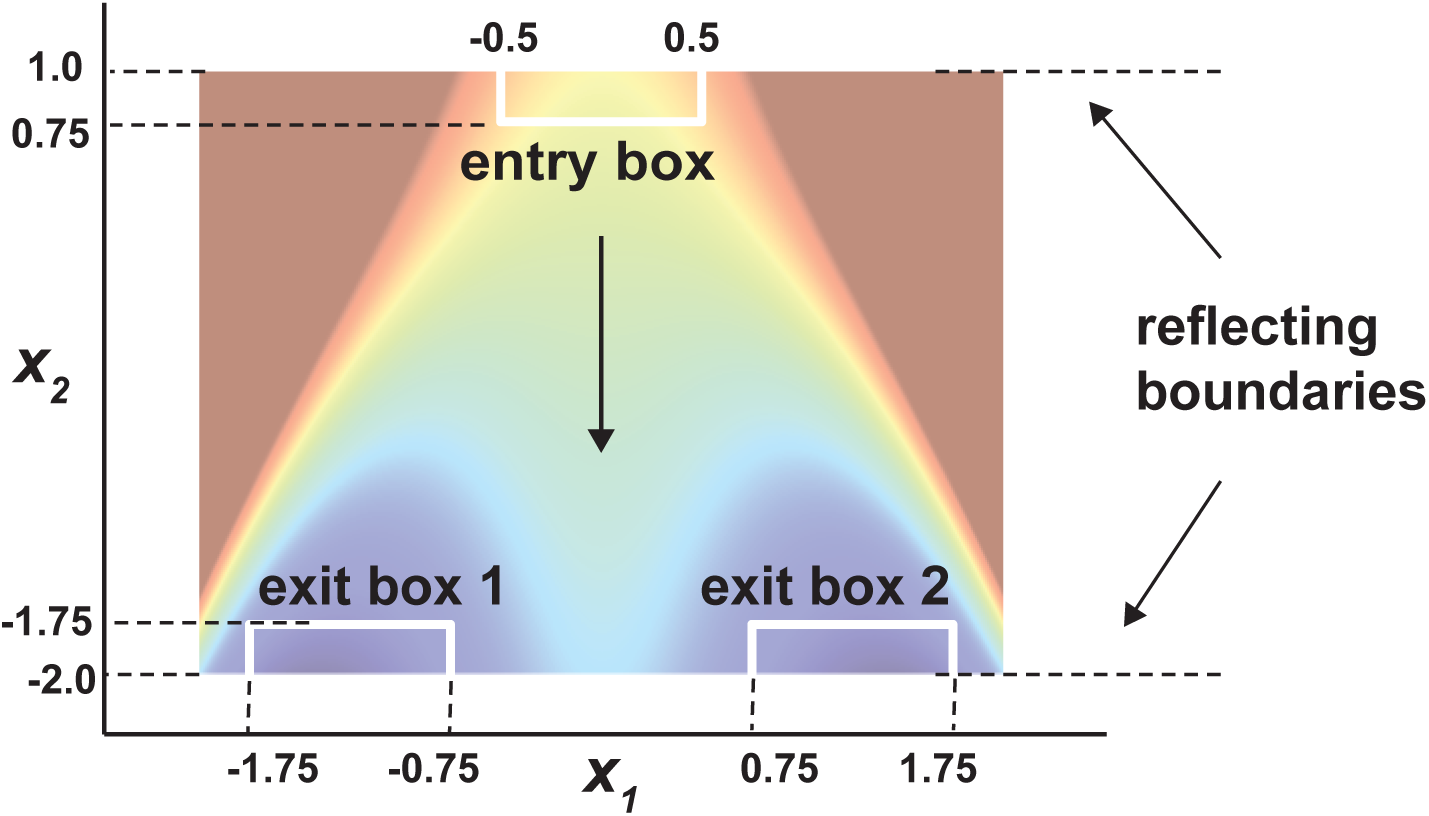
Entry/exit boundaries for a simulation of lineage bifurcation. Figure supporting Fig. 3 and Methods section 2, showing the entry/exit locations in the first two dimensions of the m-dimensional gene expression simulation. Each cell was generated at simulation time t=0 in a gene expression state chosen uniformly at random in the indicated m-dimensional source box. Cells exit through one of the two sink boxes.

**Supplementary Figure 2:**
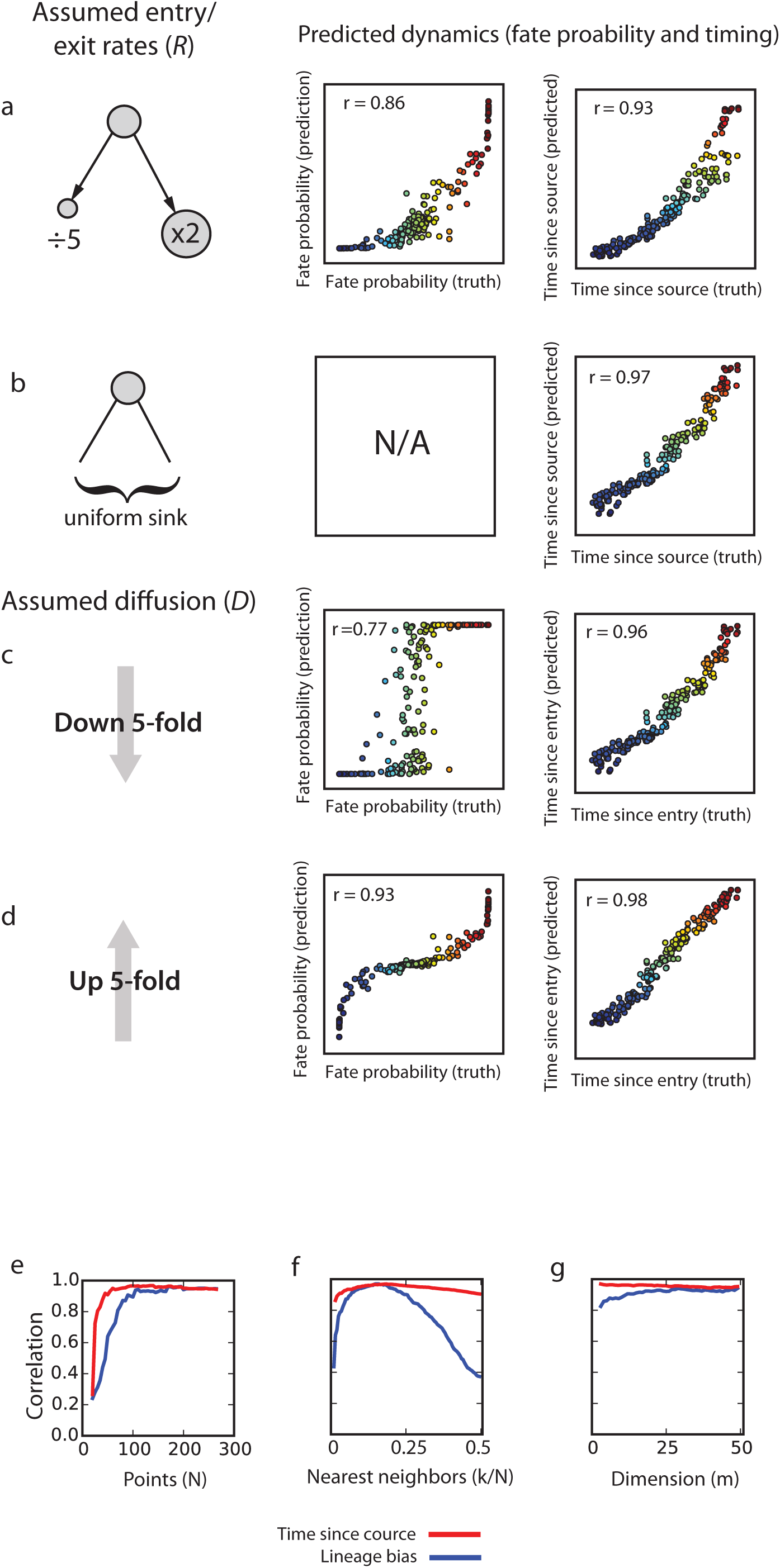
Robustness tests for the PBA algorithm. (a-d), Comparison of PBA predictions to “true” (simulated) fate bias and temporal order under imprecise assumptions about the entry/exit and diffusion parameters, D and R. These analyses also showed that: (a) imprecis,ely estimating the exit rates between two fates with a tenfold error skews estimated fate probabilities but maintains high correlations; (b), treating every point as an exit does not diminish the accuracy of predicted temporal ordering; (c),decreasing the assumed diffusion rate predicts fate commitment to occur prematurely, causing PBA to under-estimate the number of bi-potent cells; (d), increasing the assumed diffusion rate has the opposing effect, leading to over-estimate the number of bi-potent cells. (a-c), Pearson correlation between “true” and “predicted” values of fate bias and temporal order for a range of algorithm parameter values: (e), the number of cells sampled; (f), number of graph neighbors *k* (measured as fraction of total graph size); (g), simulation dimensionality *m* (i.e. number of independent genes per cell). For each case, the relevant parameter is varied while keeping the other parameters fixed (*N*=200, *k*=20, *m*=50). In general, inference of temporal order is more accurate than fate probability.

**Supplementary Figure 3:**
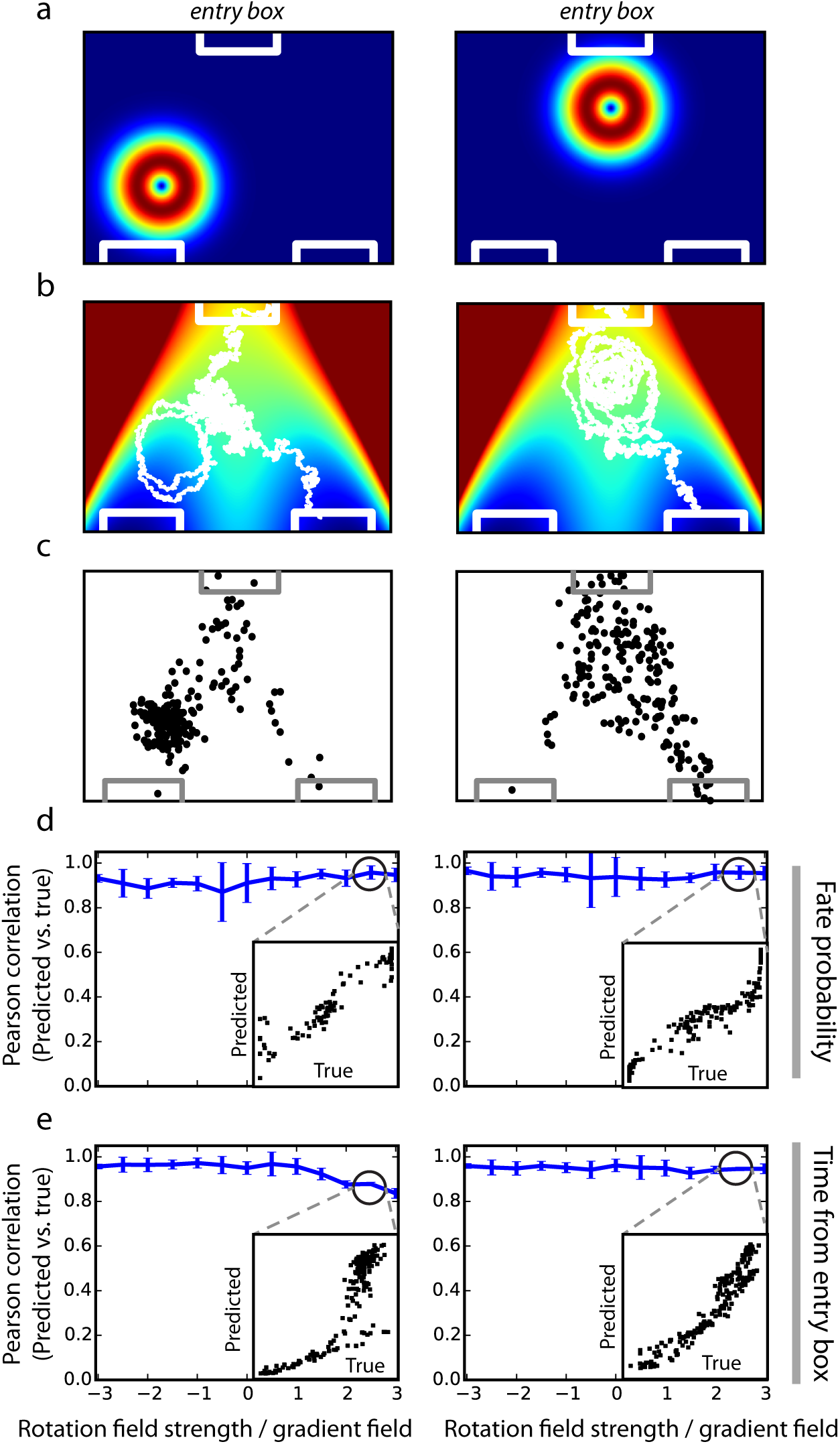
Testing PBA robustness to gene expression oscillations. PBA models gene expression dynamics as a diffusion-drift process down a potential landscape. This model makes an implicit assumption that no oscillations exist, since potential fields are irrotational. We measured the error that could be introduced by this assumption, by implanting a rotational gene expression field into the simulated fate bifurcation at two different points (left and right column), shown in (a). (b), Example simulated cell trajectories in presence of a rotational field; (c), location of sampled cells in the first two simulated gene expression dimensions; (d-e), Fidelity of PBA dynamic predictions measured by the Pearson correlation between “true” (simulated) and predicted quantities. Despite violating the assumptions of PBA, the oscillations did not significantly impact accuracy for fate probability (d) and timing (e). Blue curves indicate the mean correlation value as a function of the rotational field strength measured relative to the gradient field strength. Error bars indicate 90% confidence intervals from 10 independent trials. Panel insets show comparisons at the indicated rotational field strengths.

**Supplementary Figure 4:**
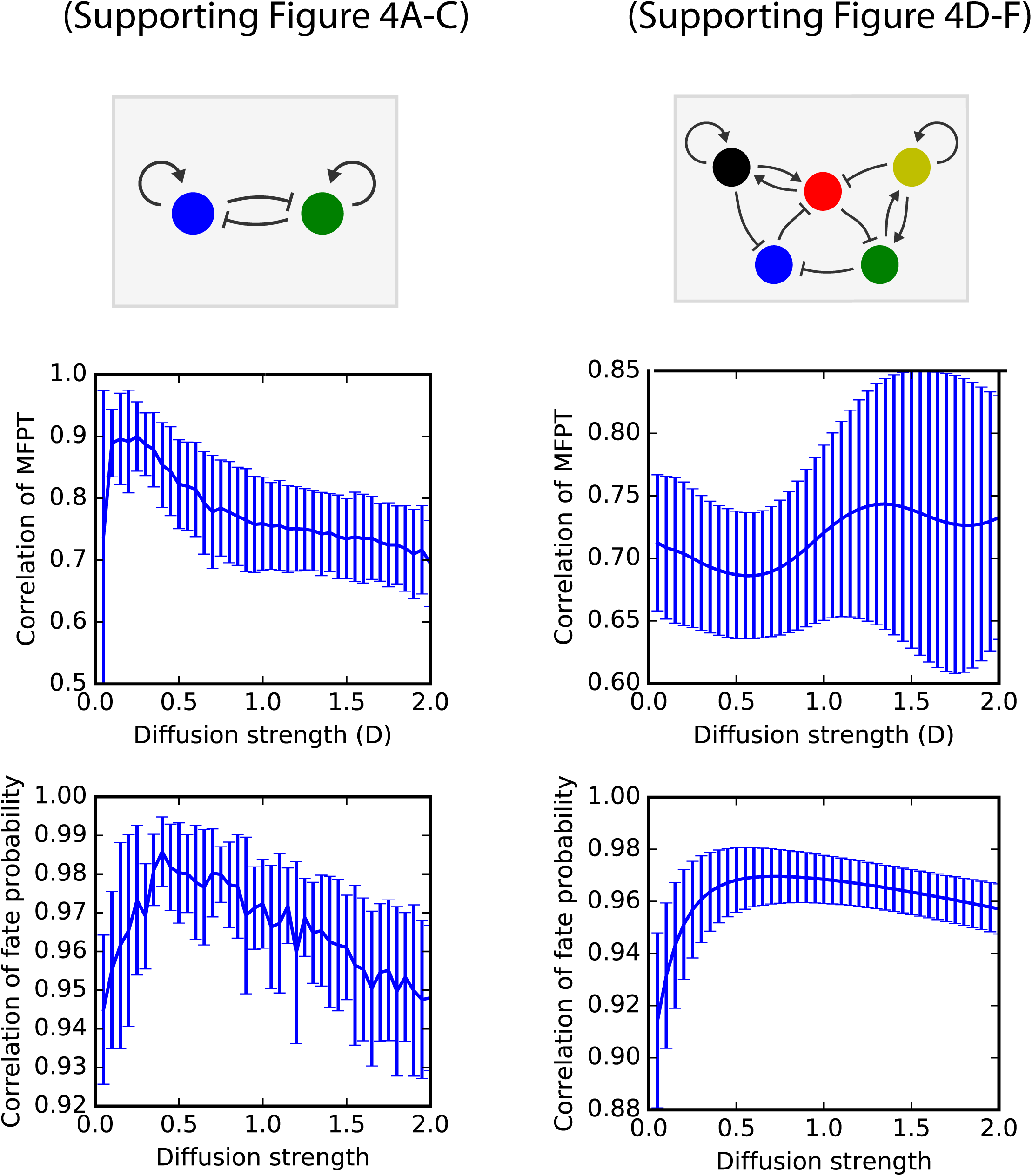
Accuracy of PBA for a range of diffusion strengths in the GRN simulations shown in Figure 4. The optimal diffusion parameter value was used in Figure 4.

**Supplementary Figure 5:**
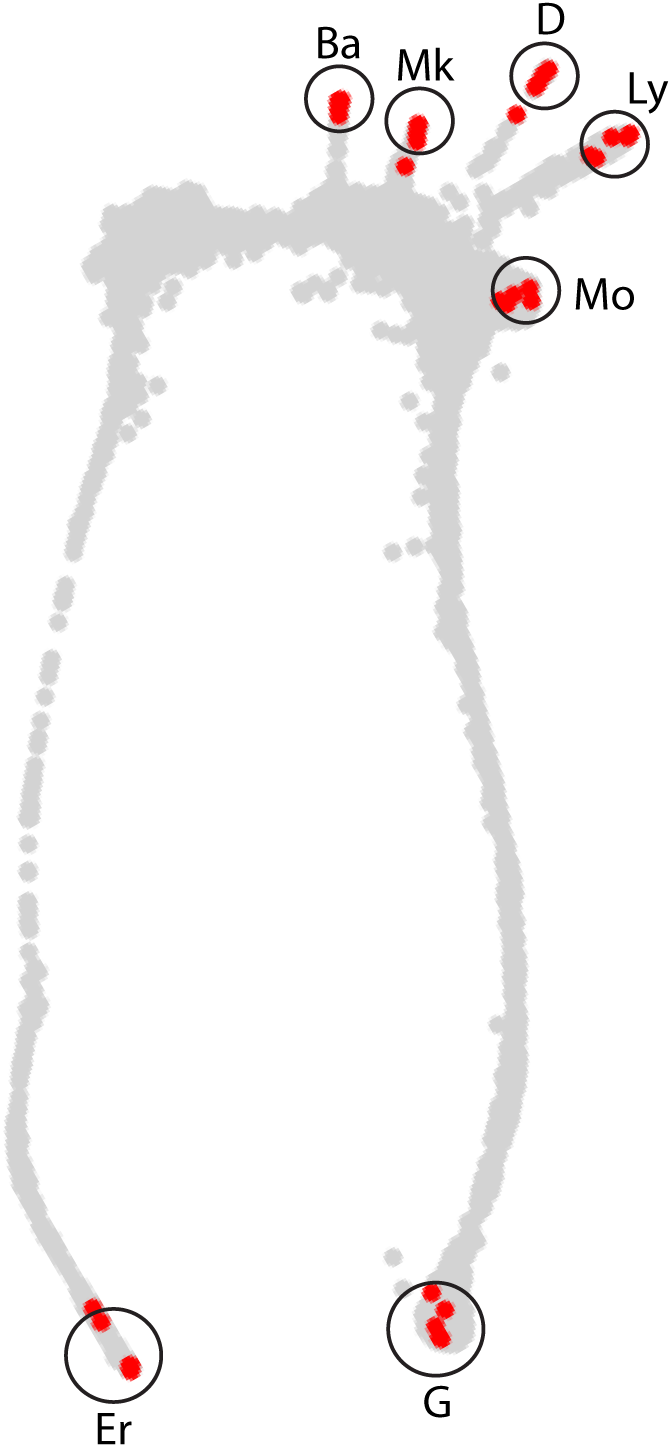
Matching cell subsets to known progenitor subpopulations. Identification of the endpoints of each lineage represented in our dataset, which occur when Kit is down-regulated. For each of seven fates, we identified endpoints as the 10 cells (red dots) with highest standardized (z-score) expression of known marker genes (Supplementary Table 1).

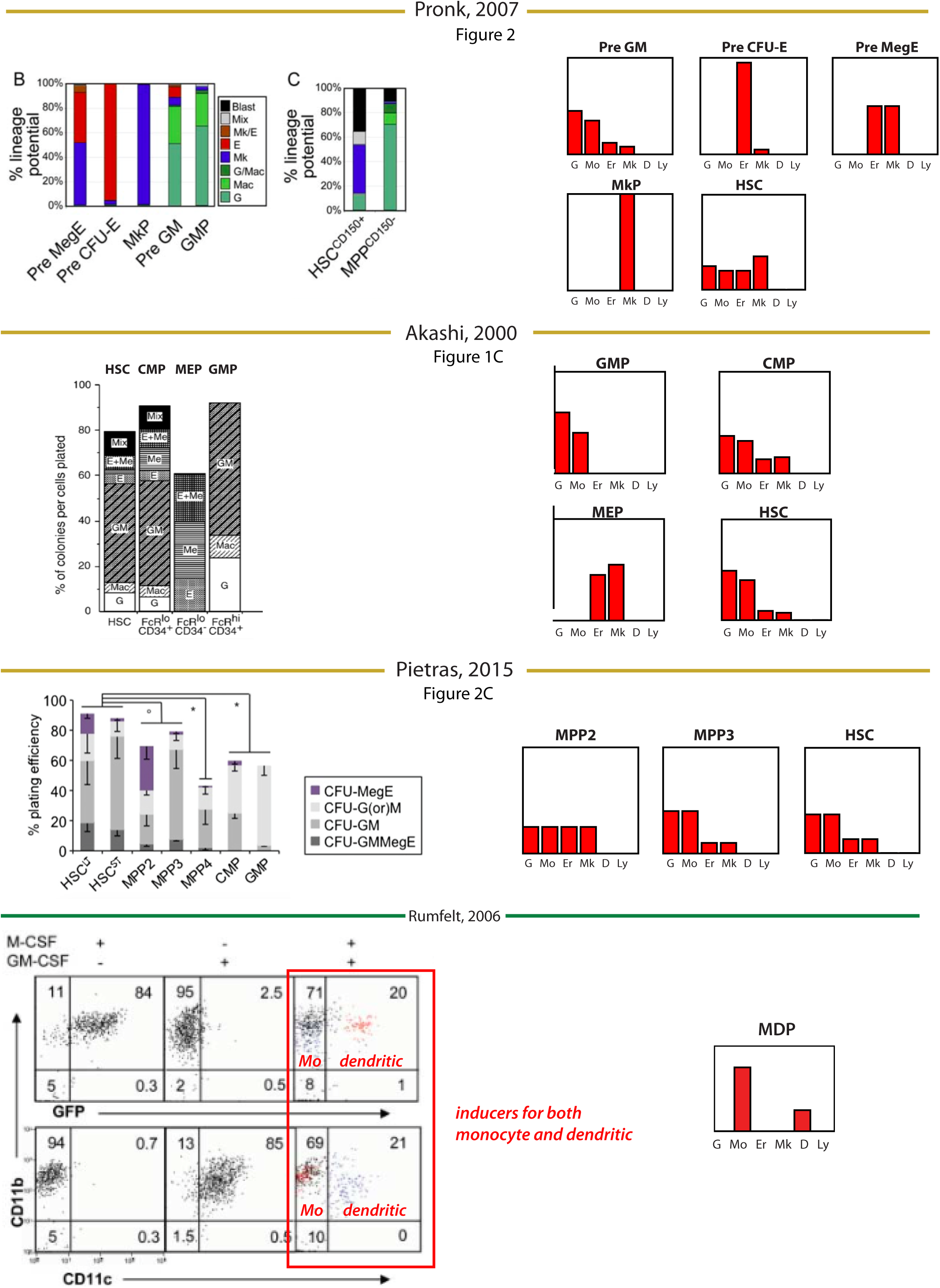

**Supplementary Figure 7:**
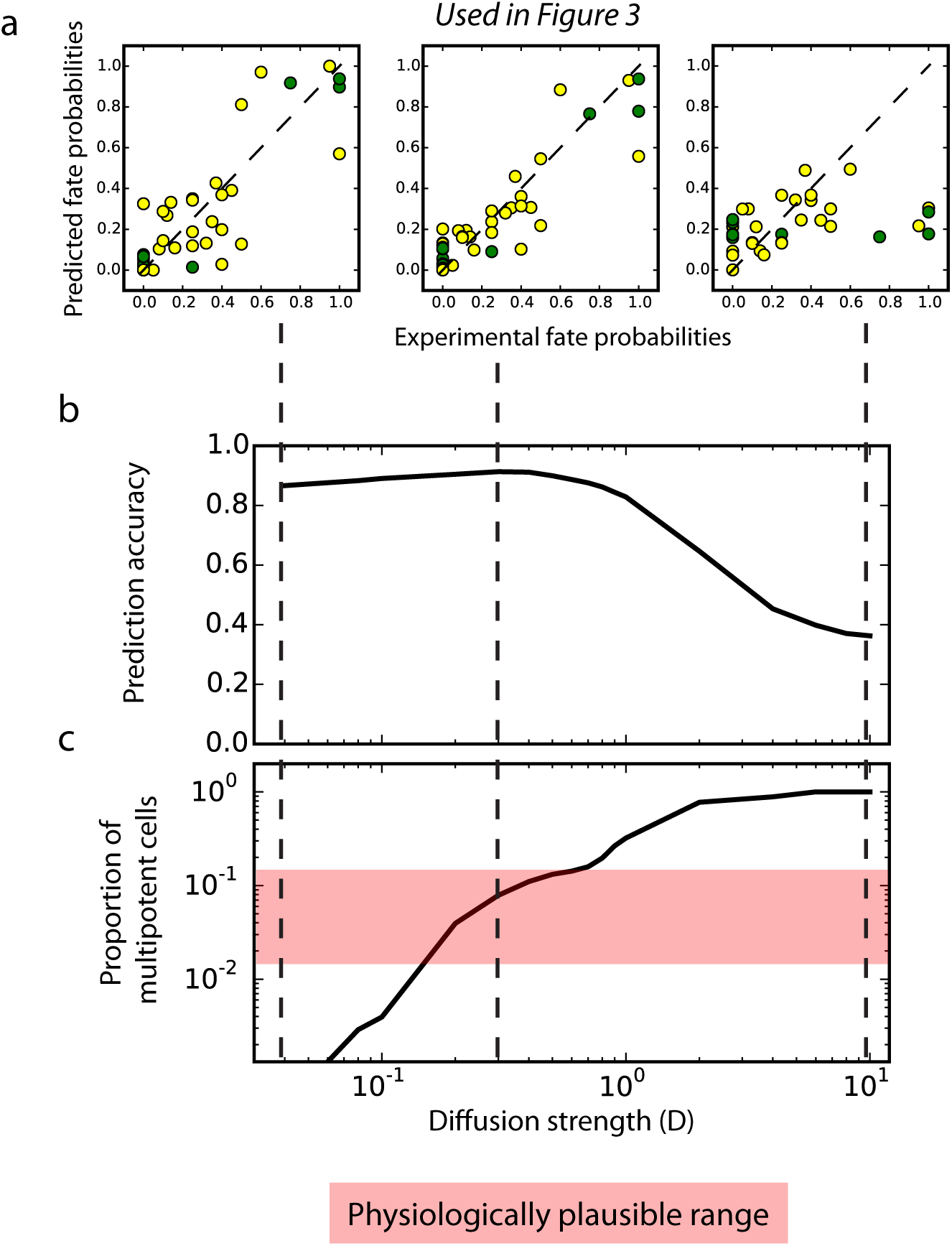
The PBA diffusion parameter (D) is constrained by subpopulation fate probabilities and by the fraction of multipotent cells. (a), The diffusion constant (D) sets the stochasticity of the PBA model and impacts the predicted fate probabilities for HPCs. (b), A systematic scan of *D* values shows that prediction accuracy remains high over a broad range of *D* values. (c), The PBA-predicted proportion of multipotent cells plotted as a function of *D.* The physiological range is highlighted (pink) (see Methods). This analysis reveals a narrow range of physiologically plausible D values that includes the point of maximum prediction accuracy. Dashed lines relate the panels in (a) to values of *D* plotted in (b,c).

